# Chromatin priming and co-factor availability shape lineage response to the neuronal pioneer factor ASCL1 in pluripotency

**DOI:** 10.64898/2026.03.19.712999

**Authors:** Jethro Lundie-Brown, Rosalind Drummond, John-Poul Ng-Blichfeldt, Roberta Azzarelli, Anna Philpott

## Abstract

Transcription factors act within defined developmental windows, yet how naïve pluripotent cells acquire competence to execute specific transcription factor-driven fate programmes remains unclear. Pioneer transcription factors that engage target sites in closed chromatin to initiate gene expression programmes often act at the top of hierarchies in cell identity transitions. However, we show that the ability of ASCL1 to induce a coherent neuronal programme emerges only after exit from pluripotency, coincident with progressive chromatin remodelling and accumulation of permissive histone marks at neuronal ASCL1 target sites. Binding analysis reveals that although ASCL1 can access a subset of neuronal loci in mESCs and EpiLCs, ASCL1 is preferentially diverted to non-neuronal sites, resulting in divergent transcriptional responses. Increasing global histone acetylation enhances activation of individual neuronal genes but is insufficient to drive full neuronal differentiation. In contrast, co-expression of the homeodomain transcription factor PHOX2B redirects ASCL1 towards neuronal targets while suppressing inappropriate programmes in mESCs. These findings demonstrate that ASCL1 pioneer activity is highly context-dependent and that developmental priming of chromatin is essential for appropriate lineage specification.

**HIGHLIGHTS:**

- Ectopic ASCL1 drives non-neuronal transcriptional responses in naïve and formative pluripotent cells
- ASCL1 occupies distinct, predominantly non-neuronal genomic targets in pluripotent cells due to differential chromatin accessibility
- ASCL1 pioneer activity is locus- and cell type-specific and predicted by histone acetylation status
- Co-expression of ASCL1 with *Phox2* homeodomain cofactors potentiates neuronal lineage acquisition in pluripotent cells

## INTRODUCTION

Development requires precise control over the generation of diverse cell types. Achieving the correct spatial and temporal coordination of growth, differentiation, and morphogenesis requires cells to regulate their competence to respond to inductive cues. However, the mechanisms by which cells acquire and maintain this competence remain poorly understood.

Mouse embryonic stem cells (mESCs) have proven an invaluable tool for studying competence for differentiation. The transition from naïve to formative pluripotency has revealed insights into the reorganisation of chromatin and transcription factor availability that prepares cells for subsequent state transitions.^1–5^ However, the effect of these transitions on competence for differentiation in response to lineage-specifying transcription factors (TFs) remains incompletely understood. ESCs are widely used in forward programming strategies, in which TF overexpression drives direct conversion to specific cell types in vitro.^6–9^ These approaches typically require prior priming through altered culture conditions^7^ or embryoid body formation,^9^ suggesting that naïve pluripotent cells are not inherently competent to respond to lineage-specifying TFs. Indeed, recent work showed that mESCs are refractory to directed differentiation induced by ASCL1,^10^ but a mechanistic understanding of this phenomenon remains elusive. Understanding how such TFs act in naïve pluripotent cells would therefore clarify both the biology of this foundational cell state and the epigenetic basis by which competence for TF-driven cell fate change is established.

Competence to respond to TFs can be shaped by chromatin patterning.^11^ Chromatin in naïve pluripotency is distinct from other tissues and reflects their capacity for multilineage differentiation and their transient nature in development.^12–15^ Naïve pluripotent cells typically have higher levels of chromatin accessibility compared with differentiated cells.^12,16,17^ Although open chromatin does not necessarily determine enhancer activity,^16^ active enhancers usually exhibit local depletion of nucleosome occupancy.^18^ Many enhancers in early development are accessible but maintained in a poised state marked by the presence of both activation-associated H3K4me1 and repressive H3K27me3, rather than the H3K27ac signature of active enhancers.^18–21^ The ability of a cell to interpret and respond to lineage-specifying TFs is therefore determined partially by the availability of TFs and their recruitment to these regulatory elements, and partially by the chromatin states that permit or restrict their binding and activity.^22–24^

Certain TFs, however, have the ability to bind their preferred motifs in nucleosome-dense chromatin and initiate opening and activation of target gene expression.^25,26^ These so-called pioneer factors often sit at the apex of transcriptional networks to drive an entire lineage program during both development and reprogramming,^27,28^ in some cases directly establishing competence for differentiation.^25,29,30^ The extent to which these pioneer TFs are subject to the same restrictions imposed by the chromatin landscape described above is not clear. Moreover, the simple distinction between pioneer and non-pioneer TFs has been challenged,^31^ as some pioneer factors display different preferences for open and closed chromatin across cell types and at different loci.^32–35^ Their activity is also frequently modulated by cofactors that refine or redirect their function, and TFs often cooperate to control cell fate in development and reprogramming.^11,36–39^

ASCL1, a proneural bHLH TF, drives both fibroblast-to-neuron conversion^40,41^ and neural stem cell differentiation *in vivo* and *in vitro*.^42,43^ ASCL1 has been described as a pioneer factor due to its ability to bind nucleosomal DNA directly.^44^ However, it also binds partially accessible chromatin and recruits chromatin remodelling enzymes.^35^ Evidence from somatic reprogramming has further suggested that competence for ASCL1-mediated conversion correlates with a specific epigenetic signature at its binding sites, ^40,41^ yet how epigenetic constraints shape the developmental acquisition of this competence remains largely unexplored. ASCL1-mediated differentiation therefore represents a tractable system for dissecting how the epigenetic landscape of early developmental states determines competence to respond to lineage-specifying pioneer TFs.^7,10,45,46^

Here, we investigate when and how cells acquire competence to respond to ASCL1 during the exit from pluripotency. By integrating RNA-seq, ASCL1 ChIP-seq, and chromatin accessibility mapping across naïve mESCs, epiblast-like cells (EpiLCs), and neuroectoderm (NE) states, we show that ASCL1 binding is highly context-dependent: key neuronal target loci are inaccessible and transcriptionally refractory in pluripotent cells, while genes associated with divergent pathways are readily activated. We find that the onset of competence to respond to ASCL1 with a coherent neuronal programme correlates with progressive chromatin remodelling and acquisition of permissive histone modifications at neuronal ASCL1 target sites. The ability of ASCL1 to drive neurogenesis is therefore dependent on developmental priming of chromatin that is established after the exit from pluripotency. We further show that the barrier to activation in mESCs for neuronal ASCL1 target genes can be partially overcome by elevating levels of histone acetylation, or co-expressing homeodomain TFs that bias ASCL1 activity towards neuronal targets. Together, these findings establish that ASCL1 pioneer factor activity is constrained in an early developmental context and reframe our understanding of how competence for somatic differentiation in response to transcriptional cues is encoded in the epigenome.

## RESULTS

### An in vitro model of early development with inducible ASCL1 activity reveals changes in competence for neuronal differentiation

Our previous work has shown that naïve mouse embryonic stem cells (mESCs) do not respond to ASCL1 overexpression by activating a neurogenic programme.^10^ Here we set out to understand why ASCL1 fails to drive neuronal differentiation in mESCs and investigate what, if any, is the transcriptional response to ASCL1 in mESCs and epiblast-like cells (EpiLC) compared to the response in neuroectodermal (NE) cells. To this end, we developed an mESC-based system with inducible ASCL1 activation (iASCL1 mESCs). HA-tagged ASCL1 was fused with the ligand-binding domain of the glucocorticoid receptor (LBD-GR),^43,47,48^ enabling conditional control by the addition of dexamethasone (Dex) to the culture medium (Fig. 1A and Fig. S1B).^49,50^ The inducible ASCL1-GR transgene (iASCL1), driven by a constitutively active promoter, was inserted at the *Rosa26* locus (see Methods) (Fig. 1A). Constitutive expression of iASCL1 was confirmed in iASCL1 mESCs (Fig. S1C). iASCL1 mESCs can be propagated in standard 2iLIF pluripotency conditions,^51^ where cells express pluripotency markers, NANOG and OCT4, in the absence of Dex (Fig. S1A).

**Figure 1.**
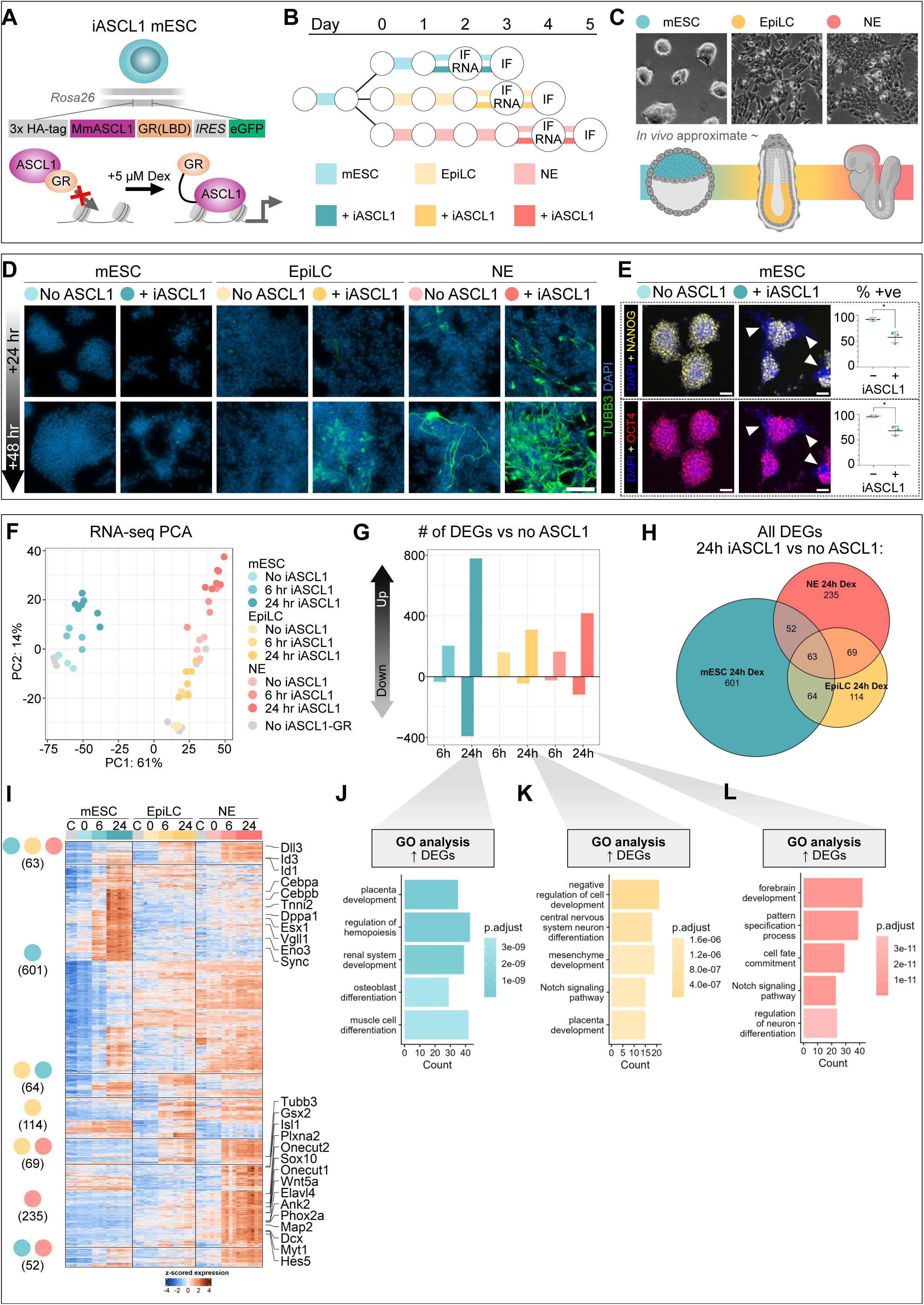
Inducible ASCL1 drives divergent and non-neuronal transcriptional programmes in mESC and EpiLC. A) Schematic of mouse embryonic stem cells with inducible ASCL1 (iASCL1 mESC). B) Differentiation timeline and sample collection for IF (D–E) and RNA-seq (F–L). “+ ASCL1” indicates 5 µM dexamethasone treatment. C) Phase-contrast images of mESC, EpiLC, and NE at day 2 without ASCL1, with schematic of in vivo approximates. D) IF showing mESC, EpiLC, and NE ± iASCL1 for 24 h (top) or 48 h (bottom). TUBB3 (green) marks neuronal differentiation. Scale bar = 100 µm. Representative of 4 experiments. E) IF of iASCL1 mESC ± 24 h iASCL1 stained for NANOG (yellow) and OCT4 (red) and percentage of positive cells (n=3; 8–10 images per experiment). Scale bar = 100 µm. * p < 0.05, t-test. F) PCA of top 1000 variable genes across conditions; grey points denote Dex-treated control line with no transgene. G) Number of differentially expressed genes (DEGs) at 6 h and 24 h relative to 0 h in each cell type (mESC blue, EpiLC yellow, NE pink; positive upregulated, negative downregulated). H) Venn diagram of overlap between ASCL1-upregulated DEGs across cell types after 24 h iASCL1. I) Heatmap of z-scored expression for DEGs upregulated by ASCL1 in at least one cell type; clusters grouped by cell-type specificity with selected genes indicated. J–L) Top 5 GO terms for upregulated genes after 24 h ASCL1 in mESC (J), EpiLC (K), and NE (L).

Culture of mESCs in N2B27 media without 2iLIF spontaneously gives rise to anterior neural ectoderm cells (NE) (Fig. S1D).^52,53^ Expression of SOX1, an early marker of NE,^54,55^ was detected at low levels from days 2 and 3 and at high levels throughout cultured cells from day 4 (Fig. S1D–E). This was concomitant with a loss of pluripotency marker OCT4 between days 3 and 4 (Fig. S1D) and followed by PAX6 and sparse TUBB3 expression from day 5, indicating spontaneous but limited neuronal differentiation from NE by this later stage (Fig. S1D). NE cells at this stage do not express endogenous ASCL1.^56^ mESC to NE differentiation proceeded via a NANOG-negative, OTX2-positive state resembling EpiLCs in the transition from naïve to formative pluripotency,^5,57^ but this identity was transient and was lost by day 2 (Fig. S1E). The EpiLC identity can be stabilised with activin A and FGF2,^3,5^ enabling interrogation of the formative state without spontaneous differentiation (Fig.1B–C and Fig.S1F). Together, mESCs, EpiLCs and NE cells constitute an *in vitro* model of the transition from pluripotency through to neuroectoderm after gastrulation (Fig.1B & C).

We next characterised the response to iASCL1 across these cell types. In NE cells, activation of iASCL1 produced limited TUBB3-positive cells within 24 hours (Fig. 1D) but by 48 hours, neuronal differentiation was widespread and TUBB3 expression was robust compared to the sparse, spontaneous neuronal differentiation observed in NE cells without activated iASCL1 (Fig. 1D). EpiLCs displayed sparse TUBB3 expression only after 48 hours of activated iASCL1, with morphology that indicates less advanced neuronal differentiation (Fig. 1D). As expected,^10^ mESCs failed to upregulate TUBB3 with or without iASCL1 over this period (Fig. 1D). This confirmed that competence for differentiation in response to ASCL1 activity is acquired during the exit from pluripotency. To exclude the possibility that resistance to iASCL1 in mESCs is attributable simply to inhibitors present in the media, we also assessed the response to iASCL1 in variants of 2iLIF lacking either the ERK inhibitor (Chiron-LIF) or the GSK3β inhibitor (PD-LIF), as well as in serum-LIF which also maintains pluripotency of mESCs^58–60^ (Fig. S1G). ASCL1-driven expression of target genes (*Tubb3* and *Hes5*) was observed exclusively in NE cells and not in these variant 2iLIF conditions, indicating that neither ERK inhibition nor GSK3β inhibition alone was responsible for suppressing neuronal differentiation (Fig. S1G).

During ASCL1-mediated differentiation, neural progenitor cells undergo rapid cell cycle exit^61–63^ and dismantle the progenitor regulatory network.^61–64^ We asked whether pluripotent cells responded in a similar way, despite the failure to induce neuronal differentiation. We found that 24 hours of iASCL1 activation in 2iLIF mESCs resulted in a significant reduction in the number of NANOG- and OCT4-positive mESCs (from 90–100% to 55–60%, p < 0.05) (Fig. 1E) and a decrease in the number of EdU-positive cycling cells (from 65% to 40%) (Fig. S1H). This is consistent with previous reports of ASCL1 activity accelerating loss of pluripotency genes in differentiating mESCs.^7^ These results indicate biological activity of iASCL1 in mESCs and suggest that the failure of mESCs to differentiate into neurons in response to ASCL1 is not caused by intrinsic resistance to ASCL1-mediated transcriptional activation.

### ASCL1 activates distinct transcriptional profiles in pluripotent, epiblast-like and neuroectodermal cells

To investigate the transcriptional response to iASCL1 genome-wide in mESC, EpiLC and NE, we performed RNA-seq in cells exposed to 0, 6 or 24 hours of iASCL1 activity before collection (Fig. 1B and 1F). ASCL1 altered the expression of numerous genes in all three cell types (Fig. 1G and Fig. S1I–K) (Table S1) demonstrating its transcriptional activity in all three contexts. Moreover, the greatest number of upregulated differentially expressed genes (DEGs) after 24 hours of ASCL1 activation was seen in mESCs (mESC: 780; EpiLC: 310; NE: 419) (Fig. 1G). Fewer genes were downregulated than upregulated in all cell types (Fig. 1G), consistent with ASCL1’s established predominant role as an activator rather than a repressor.^43^ Subsequent analysis therefore focused on upregulated DEGs. A set of 63 “universal” ASCL1-responsive DEGs upregulated in all three contexts was identified, including known ASCL1- and Notch-related targets, *Dll3*, *Id1*, and *Id3*. However, the majority of ASCL1-upregulated DEGs after 24 hours were cell type-specific, including 235 genes that were only upregulated by ASCL1 in NE, 114 in EpiLC and 601 in mESC (Fig. 1H–I).

To better understand the functional identity of genes activated by ASCL1 in each cell type, we performed gene ontology (GO) analysis on upregulated DEGs from mESCs, EpiLCs and NE cells (Fig. 1J–L) (Table S2). iASCL1-responsive DEGs in NE were associated with neurogenesis-related terms including “forebrain development” and “regulation of neuron differentiation” (Fig. 1L), whereas ASCL1-responsive genes in mESC and EpiLC, were associated with terms including “placenta development”, “hemopoiesis”, “muscle cell differentiation”, and “mesenchyme development” (Fig. 1J–K), indicating activation of divergent programmes not associated with neurogenesis. Notably, ASCL1-activated genes associated with “central nervous system neuron differentiation” were upregulated in EpiLCs, yet these cells displayed only limited TUBB3-expression (Fig. 1D and 1K). This indicates that at least some of the 235 NE-only ASCL1 responsive genes must be essential for neuronal differentiation; their resistance to activation in response to ASCL1 in mESC and EpiLC likely contributes to the limited competence of these cells to undergo ASCL1-mediated neuronal differentiation.

The list of DEGs identified after 24 hours of iASCL1 activity will also include targets of secondary effectors in addition to genes that are directly targeted by ASCL1. It is possible that ASCL1 directly activates the same target genes in all cell types, and subsequent transcriptional responses diverge due to downstream effects. One advantage of our iASCL1 system is the ability to capture early transcriptional changes; we therefore also identified ASCL1-responsive DEGs after 6 hours of iASCL1 activation in each cell type (Fig. S1L–O) (Table S1). As expected, fewer DEGs were identified after 6 hours than at 24 hours, but ASCL1-responsive DEGs were already cell type-specific at this earlier timepoint (Fig. S1L) and GO analysis of these 6-hour DEGs also revealed that neuronal-associated terms were only highly represented in NE (Fig. S1M–O) (Table S2). Together, these data support a model for distinct ASCL1 activity in these three cell types that is largely shaped by differences in the initial cellular context, rather than as a result of downstream divergent responses to ASCL1 activation.

### ASCL1 binding is cell-type specific and explains divergent transcriptional responses

To activate its target genes, ASCL1 must bind to *cis*-regulatory elements (CREs) and initiate transcription. ASCL1 has been described as an on-target pioneer TF, and previous reports have claimed that its binding sites are similar in both developmental and reprogramming contexts.^40,41^ We therefore asked whether ASCL1 binds distinct loci in mESC, EpiLC and NE and whether this explains the divergent transcriptional responses we observed. We used ChIP-seq to probe genome-wide ASCL1 binding (Fig. S2A–C) and focused our analysis on the top 10,000 peaks from each cell type (Fig. S2D). This resulted in a total consensus set of 17,210 unique ASCL1 binding sites bound in at least one of mESC, EpiLC or NE (Fig.2A). Grouping these peaks revealed striking cell type-specific binding patterns for ASCL1 (Fig. 2A) (Table S3). EpiLCs and NE have more ASCL1-binding sites in common (3,181) than either mESC and EpiLC (1,353) or mESC and NE (402). A further 3,972 ASCL1-binding sites are shared in all three cell types (Fig. 2A). Distinguishing ASCL1 binding sites at promoters, introns, or distal intergenic regions revealed similar proportions in mESCs, EpiLCs and NE (Fig. S2E). Motif enrichment analysis across all ASCL1 site clusters confirmed that binding sites in all three cell types share a similar but not identical central E-box motif (CAGCTG), rather than harbouring cell type-specific E-box variants (Fig. S2F). Preferential binding of bHLH TFs to E-box variants can be altered by formation of distinct dimers,^65^ but the similarity of motifs discovered in our ASCL1 ChIP-seq suggests that this is not a primary determinant of cell type-specific ASCL1 activity in mESC, EpiLC and NE.

**Figure 2.**
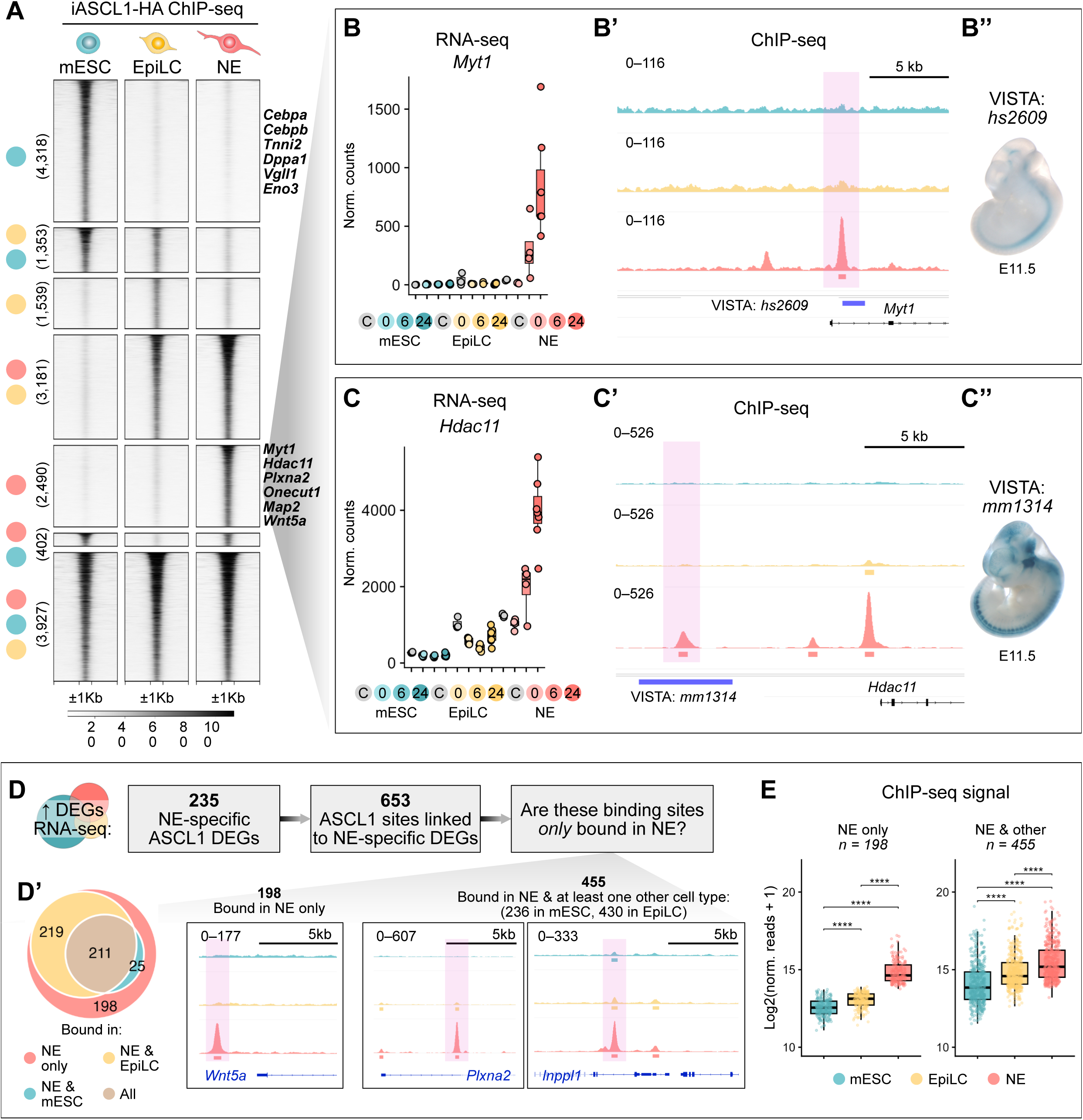
Context-dependent ASCL1 genomic occupancy underlies divergent transcriptional responses across developmental states. A) Top 10,000 ASCL1 binding sites from anti-HA tag ChIP-seq from each cell type (mESC, EpiLC, NE) collated to give 17,210 unique binding sites. ASCL1 binding sites are clustered by cell type-specific binding patterns. Mean RLE-normalised ChIP-seq signal from 3 or 4 replicates. B–C) Examples of NE-specific target genes from RNA-seq (B and C) and NE-specific ASCL1 binding sites (B’ and C’), which overlap with validated enhancers from the VISTA dataset (B’’ and C’’). *In situ* hybridisation from VISTA database shows enhancer activity in neural tube and brain at E11.5. D) Categorising ASCL1 binding sites linked to NE-specific genes based on whether they are exclusively bound in NE (left, n = 198) or bound in at least one other cell type (right, n = 455). (D’) Venn diagram for these categories. Example IGV tracks of ASCL1 ChIP-seq for each category: *Wnt5a*, *Plxna2*, *Inppl1*. E) ChIP-seq signal at sites in each of the categories in (D). One-way ANOVA with repeated measures, Benjamini-Hochberg corrected pairwise comparisons: **** p < 0.0001.

We next asked whether these divergent binding profiles of ASCL1 could explain the divergent transcriptional profiles. We therefore integrated our ChIP-seq and RNA-seq data to determine the relationship between ASCL1 binding and ASCL1-responsive differentially expressed genes, identifying all ASCL1 binding sites within 100 kb upstream or downstream of gene transcription start sites (Table S3).^66,67^ We found that a large majority of the most upregulated ASCL1-responsive DEGs in each cell type were proximal to at least one ASCL1 binding site (mESC: 685/780; EpiLC: 291/310; NE: 385/419) (Fig. S2G). Furthermore, differentially expressed neuronal genes that are only expressed in NE were associated with NE-specific ASCL1 binding sites, while non-neuronal mESC-specific ASCL1-responsive DEGs (Fig. 2A, labelled genes) were associated with mESC-specific binding sites.

To confirm the *in vivo* neuronal activity of the NE-specific ASCL1 regulatory elements, we compared these ASCL1 binding sites to the location of validated enhancers from the VISTA database.^68^ Gene expression of NE-specific DEGs and their associated NE-specific ASCL1 binding sites is consistent with *in vivo* activity of enhancers associated with these sites in the developing mouse forebrain, hindbrain and spinal cord (Fig. 2B–B’’ and Fig. 2C–C’’). These data indicate that ectopic ASCL1 in NE is recruited to physiologically relevant ASCL1 binding sites, even before endogenous ASCL1 has been activated.

NE cells undergo a strong upregulation of the neuronal programme on ASCL1 activation (Fig. 1D and L). To understand more about the mechanisms that underpin ASCL1-mediated neurogenesis, we next focused more closely on the 235 NE-specific ASCL1-responsive DEGs (Fig. 1H) and found that 208 of them were associated with an ASCL1 binding site within ± 100kb. As some of these genes were linked to multiple ASCL1 binding sites, this corresponded to 653 ASCL1 binding sites proximal to the 208 bound and NE-specific target genes (Fig. 2D) (Table S4). Since ASCL1 failed to activate neurogenesis in mESCs and EpiLCs, we then asked whether these 653 binding sites were also bound by ASCL1 in mESCs or EpiLCs or not. Surprisingly, 455 of these sites associated with genes only upregulated in NE were also bound by ASCL1 in either mESC or EpiLC, while only 198 of these sites were exclusively bound by ASCL1 in NE (Fig. 2D’). However, where ASCL1 was found to be bound to specific sites in two or more cell types, average ChIP-seq signal at the shared sites was significantly higher in NE than in mESC and EpiLC (Fig. 2E). Taken together, the ChIP-seq data suggest that ASCL1 binding, although present at some of the correct neuronal target genes, is lower in mESC and EpiLC and binding at non-neuronal sites is favoured. This may underlie the reduced competence of mESC and EpiLC to respond to ASCL1 by executing a neuronal differentiation programme.

Context-dependent binding has been reported for various TFs, including pioneer factors,^69^ but ASCL1 has been described as targeting the same sites in development and reprogramming.^40^ To further explore context-dependent ASCL1 activity, we curated a range of published ASCL1 ChIP-seq datasets from *in vitro* neural stem cells (NSCs),^40,43^ *ex vivo* adult neural progenitors,^70^ mouse embryonic fibroblasts (MEFs),^40,71^ mESCs,^46^ and embryonic neural tube^72^ (Fig. S2H) and compared with our ASCL1 binding sites from mESC, EpiLC and NE. Correlation analysis comparing the similarity of ASCL1 binding sites across these datasets revealed that ASCL1 indeed displays cell type-specific occupancy of binding sites, even across cell types that undergo ASCL1-mediated neuronal differentiation (e.g. NSCs and MEFs). Looking more closely at strength of binding at cell type-specific peaks, ASCL1 binding in our ASCL1 NE ChIP-seq was higher than in mESC or EpiLC at ASCL1 binding sites that are also found in NSC and embryonic neural tube from published datasets^43,72^ (Fig. S2I). This analysis suggests that the pioneer model of ASCL1 activity where ASCL1 accesses the same sites in a variety of chromatin states across cell types does not extend to explaining binding differences across cell types in very early development.^40^ Instead, the data indicate that its binding is context-dependent and that competence for ASCL1-mediated neuronal differentiation is underpinned by binding at specific sets of neurogenic regulatory elements that is restricted in pluripotent cells.

### Cell type-specific chromatin pre-patterning correlates with distinct binding patterns

Having established that ASCL1 occupies distinct binding sites in mESCs, EpiLCs, and NE, we next investigated chromatin features that may explain these differences. Pluripotent cells are characterised by having globally low levels of repressive heterochromatin,^12,73^ so it was striking that ASCL1 fails to bind its NE-specific sites in mESCs. We sought to better understand whether chromatin accessibility might explain distinct ASCL1 binding profiles. To examine ASCL1 binding site accessibility across cell types, we performed ATAC-seq on mESCs, EpiLCs, and NE that had been exposed to either 6 or 24 hours of iASCL1 activity before collection (Fig. 3A–B and Fig. S3A–D). We also took untreated samples at the same final time point (No ASCL1) (Fig. 3A–C) to measure chromatin accessibility that is independent of ASCL1 activity. Surprisingly, given ASCL1’s affinity for nucleosomal DNA,^44^ we found that 80–87% of ASCL1 binding sites were accessible in each cell type even in the absence of iASCL1 activity (Fig. S3E–G). Moreover, differences in chromatin accessibility between cell types in the absence of ASCL1 closely mirrored the cell type-specific ASCL1 binding patterns described above (Fig. 3C). For example, mESC-only ASCL1 binding sites are more likely to be accessible in mESC than those same sites in EpiLC or NE in the absence of ASCL1 activation, with a similar pattern seen for other cell type-specific binding sites (Fig. 3C and Fig. S3H). This indicates preferential recruitment of ASCL1 to binding sites that are already accessible and that these vary between cell types.

**Figure 3.**
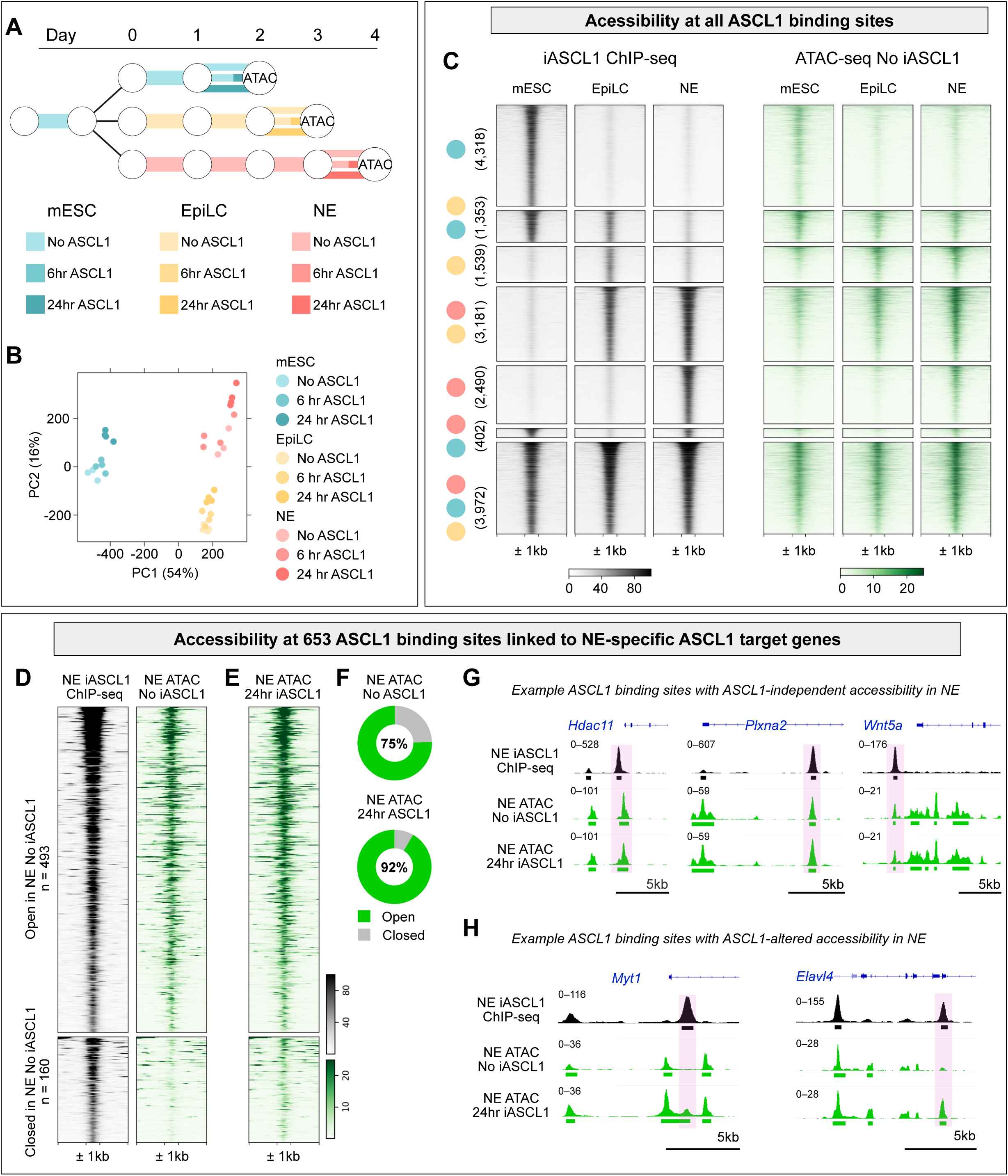
Chromatin accessibility dictates ASCL1 binding, with pioneer activity restricted to a minority of neuronal target sites. A) Schematic for ATAC-seq experiment. Samples were exposed to either no ASCL1, 6 hrs, or 24 hrs before collection at the same endpoint. B) PCA for ATAC-seq samples C) Heatmap of ASCL1 ChIP-seq (black, left) and ATAC-seq signal with No ASCL1 (green, right) clustered by cell type-specific binding patterns defined in Fig. 2A. Row order is preserved across the two heatmaps. Heatmaps centred on ASCL1 peak ± 1 kb. D) Heatmaps of ASCL1 ChIP-seq (black) and ATAC-seq (No ASCL1) (green) for 653 ASCL1 sites linked to NE-specific DEGs defined in Fig. 2D. Peaks are clustered by whether they are open (top, n = 493) or closed in NE (No ASCL1) (bottom, n = 160). Heatmaps centred on ASCL1 peak ± 1 kb. E) Heatmap of ATAC-seq (24hr ASCL1) for same 653 ASCL1 sites linked to NE-specific DEGs with row order preserved from (D). Heatmaps centred on ASCL1 peak ± 1 kb. F) Donut plots to show percentage of 653 ASCL1 binding sites in D and E that are open (green) or closed (grey) in No ASCL1 ATAC-seq (top) or 24 hr ASCL1 ATAC-seq (bottom). G) IGV tracks of ASCL1 binding sites that are open in NE (No ASCL1) ATAC-seq. H) IGV tracks of ASCL1 binding sites that are closed in NE (No ASCL1) ATAC-seq but become open in NE (24 hr ASCL1) ATAC-seq.

We focused our analysis on the previously defined 653 ASCL1 binding sites associated with NE-specific ASCL1 target genes and asked whether these sites were open or closed in NE in the absence of iASCL1 activation (Table S4). Consistent with the global pattern described above, 493 (75%) of these 653 ASCL1 binding sites were open in NE even without iASCL1 activity (Fig. 3D and 3F). Therefore, most of the binding sites linked to NE-specific target genes are open in NE independently of ASCL1 (Fig. 3G and Fig. S3I). The remaining 160 (25%) of these 653 ASCL1 binding sites were inaccessible in the absence of ASCL1 (Fig. 3D and 3F). After 24 hours of iASCL1 activity, 598 (92%) of the 653 sites were open, indicating 105 of 160 initially closed sites had been rendered accessible after 24 hours of ASCL1 activity (Fig. 3E–3F and 3H). Therefore, in NE the majority of binding sites associated with NE-specific gene expression were open independently of ASCL1 activity, while additional further sites were pioneered after ASCL1 activation (Fig. 3E–F and 3H).

Given this strong preference for ASCL1 to bind to accessible chromatin in NE, we asked whether the 653 ASCL1 binding sites linked to NE-specific gene activation were or were not accessible in mESCs. Surprisingly, we found that 55% of these sites were also accessible in mESCs in the absence of ASCL1 (Fig. 4A–B). Since the genes associated with these open sites are not responsive to ASCL1 in mESCs, chromatin accessibility alone cannot fully account for the divergent transcriptional response to ASCL1 at these loci. We therefore asked why transcriptional activation of these genes fails in mESC despite the fact that many of the ASCL1 binding sites associated with them are open and bound by ASCL1 (Fig. 4A–B). We reasoned that histone modifications might provide an additional layer of gene regulation.

**Figure 4.**
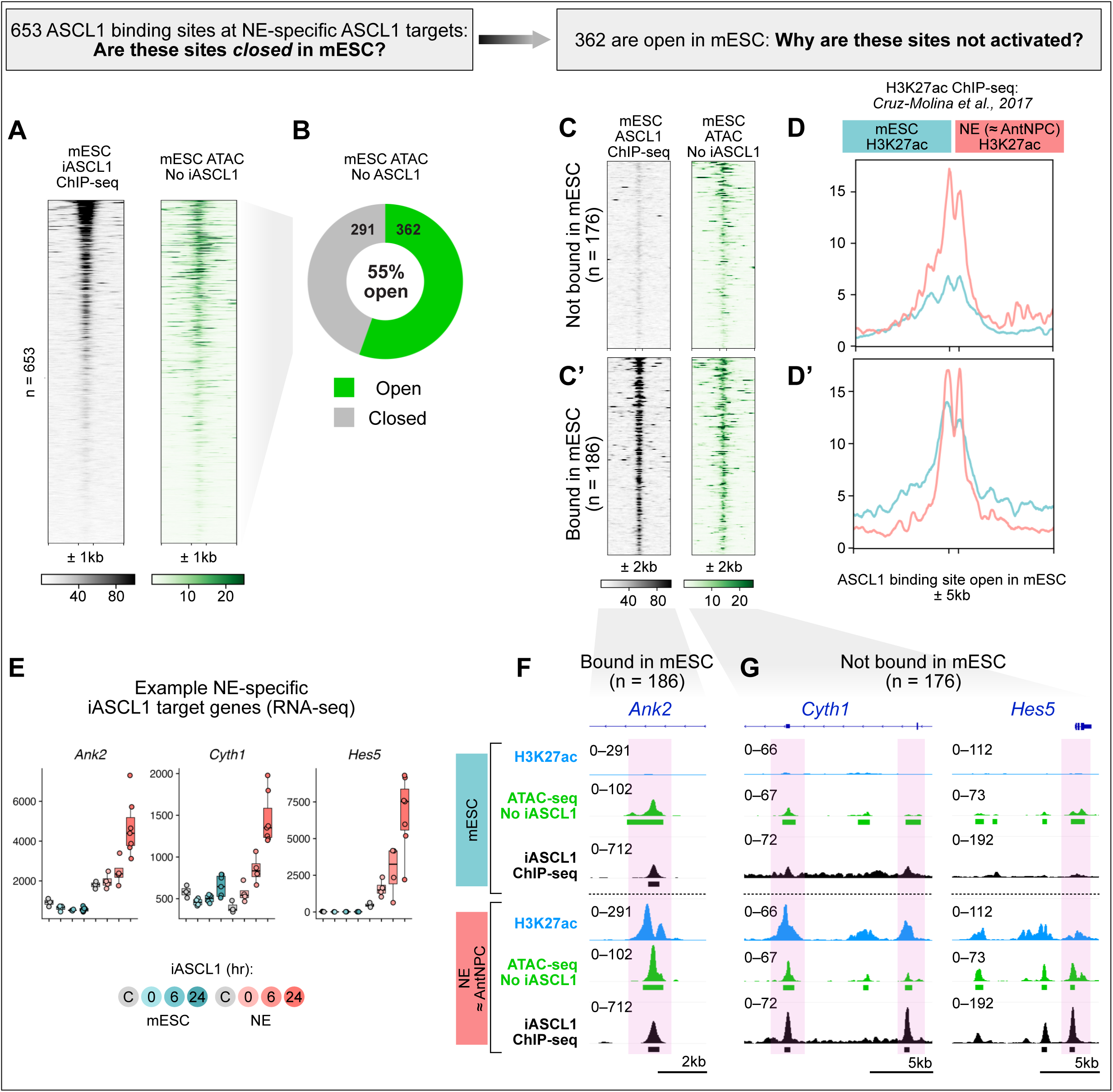
Histone acetylation status at shared accessible sites predicts productive ASCL1 binding and target gene activation. A) Heatmaps of ASCL1 ChIP-seq (black) and No ASCL1 ATAC-seq (green) from mESC at the 653 ASCL1 binding sites associated with NE-specific ASCL1 target genes defined in Fig. 3D. Heatmaps centred on ASCL1 peak ± 1 kb. B) Donut plot of percentage of the 653 sites that are open (green) or closed (grey) in mESC (No ASCL1) ATAC-seq data. C) Heatmaps of ASCL1 ChIP-seq (black) and ATAC-seq (No ASCL1) (green) from mESC for the 362 out of 653 sites that are open in mESC (No ASCL1) ATAC-seq data. Peaks are clustered based on whether are (C’) bound in mESC (bottom, n = 186) or (C) not bound in mESC (top, n = 176). Heatmaps are centred on ASCL1 peak ± 2 kb. D) H3K27ac ChIP-seq signal at 176 ASCL1 binding sites in (C); D’) H3K27ac ChIP-seq signal at 186 ASCL1 binding sites in (C’). H3K27ac data is shown for mESC (blue) and AntNPC (pink) from Cruz-Molina et al., 2017^20^. Profiles are centred around ASCL1 binding sites ± 5 kb. E) Normalised RNA-seq counts for representative NE-specific ASCL1 target genes (*Ank2*, *Cyth1*, *Hes5*) in mESC (blue) and NE (pink) exposed to 0, 6, or 24 hours of iASCL1 activity. “C” (grey) represents Dex-treated control line with no iASCL1 transgene. N = 4. F–G) IGV tracks to show ASCL1 binding sites near the NE-specific target genes in (D) that are bound (F) or not bound (G) by ASCL1 in mESCs. mESC H3K27ac (blue), mESC No ASCL1 ATAC-seq (green), mESC ASCL1 ChIP-seq (black); NE (AntNPC) H3K27ac (blue), NE No ASCL1 ATAC-seq (green), NE ASCL1 ChIP-seq (black). Rectangles below tracks indicate a called peak. Peaks are highlighted in pink boxes.

Histone H3 acetylation at lysine 27 (H3K27ac) and methylation of lysine 4 (H3K4me1) both correlate with transcriptional activity of enhancers.^74^ Moreover, mESCs have been shown to maintain enhancers associated with differentiation in a poised state, marked by accessible, H3K4me1-rich chromatin but lacking H3K27ac.^18,20^ Poised enhancers are not active, but respond rapidly at the onset of differentiation.^19,20,75^ To see whether differential enrichment of these marks at ASCL1 binding sites in mESC and NE might explain the cell type-specific patterns of ASCL1 target gene activation, we analysed published histone ChIP-seq data^20,76^ from 2iLIF mESCs, EpiLCs and anterior neural precursor cells (AntNPCs) (Fig. S4A–D). AntNPCs are approximately equivalent to NE cells and they have also not yet upregulated endogenous ASCL1 expression.^20,56^ Globally, we found that cell type-specific ASCL1 binding correlated with H3K27ac (Fig. S4B) and H3K4me1 marks (Fig. S4C), consistent with preferential recruitment of ASCL1 to active chromatin. Notably, there was no enrichment of H3K4me1 at NE-specific ASCL1 binding sites in mESC or EpiLC, indicating that these sites are not maintained in a poised state in these cells (Fig. S4C’).

We then looked specifically at the 362 ASCL1 binding sites that were open in mESCs, despite only being associated with ASCL1 transcriptional activity in NE (Fig. 4A–B). Of these 362 open ASCL1 binding sites, 176 were not bound in mESCs, despite being accessible (Fig. 4C). We saw that these sites displayed low levels of H3K27ac deposition in mESC (Fig. 4D), consistent with their maintenance in inactive chromatin. On the other hand, these sites displayed high levels of deposited H3K27ac in NE (AntNPC) (Fig. 4D). This indicates that these sites are not bound or activated by ASCL1, despite their accessibility, because they are not marked with active H3K27ac. The remaining 186 sites were bound in mESC and open in mESC, even though they also displayed NE-specific transcriptional upregulation of their associated genes in response to ASCL1 (Fig. 4C’). We saw that H3K27ac was lower at these sites in ESC than in NE (AntNPC), indicating that lower levels of pre-existing H3K27ac marks contribute to a failure of ASCL1 to activate associated NE-specific genes in ESCs, even when ASCL1 is bound at these sites in both cell types. Example NE-specific ASCL1 target genes (Fig. 4E) and H3K27ac marks at their associated binding sites (Fig. 4F–G) highlight this pattern. For example, *Ank2* is bound and accessible in mESC and NE, but displays a clear enrichment of H3K27ac at its ASCL1 binding site in NE (AntNPC) compared to mESC (Fig. 4F). By contrast, *Cyth1* and *Hes5* are examples of genes that are accessible but not bound in mESC, and these sites show low levels of H3K27ac in mESCs (Fig. 4G). These findings indicate that the NE-specific ASCL1 target genes are not activated in mESCs because they are either inaccessible to ASCL1 or accessible but not marked with active H3K27ac.

### Modulating histone acetylation enhances ASCL1 transcriptional activity, but does not facilitate neuronal differentiation in mESCs

Our epigenetic data suggest that pre-existing chromatin status, particularly histone acetylation and chromatin accessibility, is the primary determinant of ASCL1 binding and cellular competence for ASCL1-mediated neuronal gene activation. We therefore asked whether artificially enhancing histone acetylation could increase the competence of mESCs to respond to iASCL1 (Fig. 5A). We treated mESCs and NE cells with 10 nM trichostatin A (TSA), a class I/II histone deacetylase (HDAC) inhibitor that broadly elevates H3K27 acetylation, and profiled the transcriptional response to iASCL1 activation by RNA-seq (Fig. 5B–C and Fig. S5A–B).

**Figure 5.**
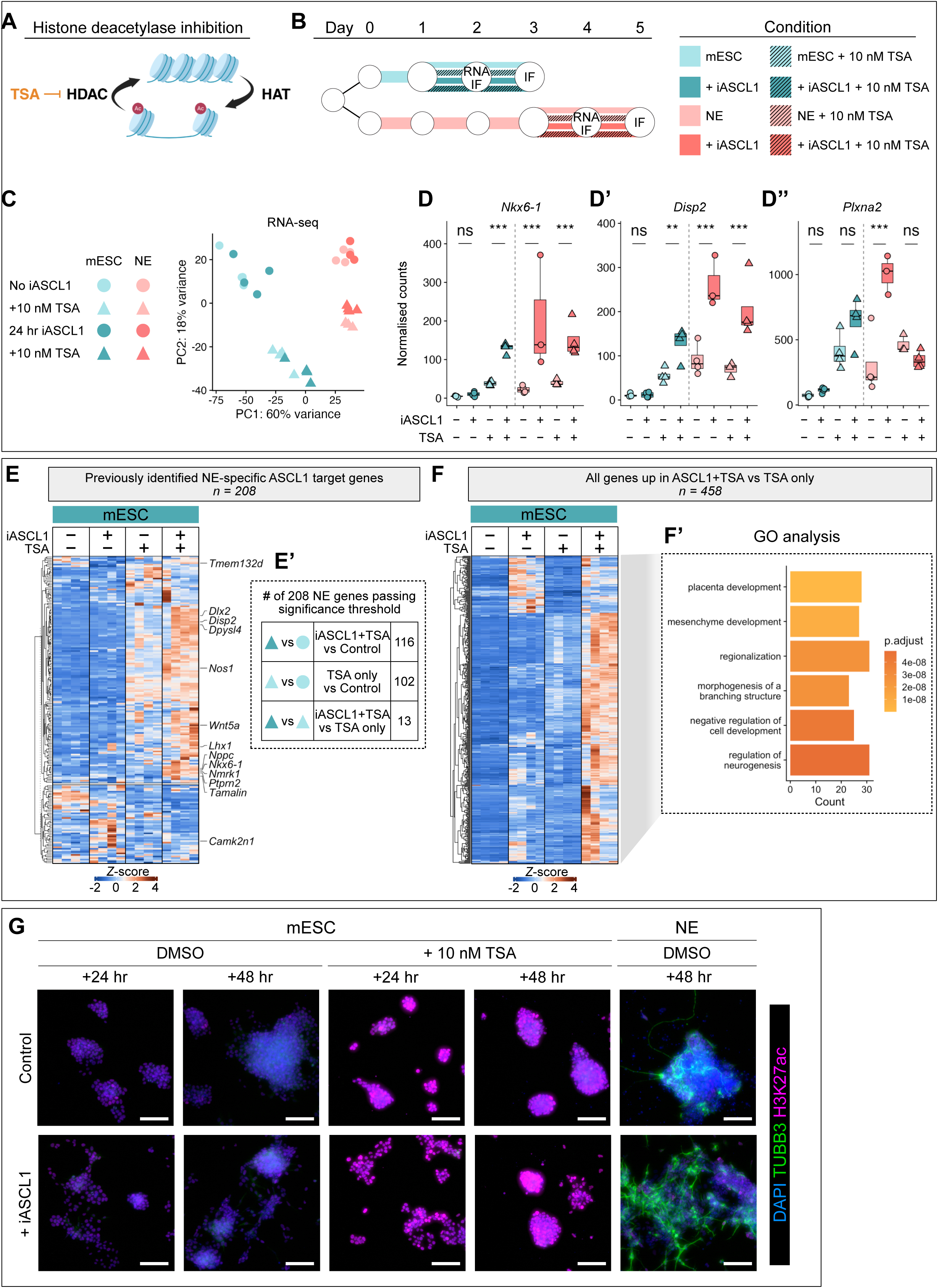
Global enhancement of histone acetylation partially unlocks neuronal gene expression by ASCL1 but fails to redirect differentiation in mESCs. A) Schematic of HDAC inhibition (HDACi) and acetylation. B) Schematic of culture and treatment of mESC (blue) and NE (pink) with iASCL1 and/or 10 nM TSA for 24 hr (RNA and IF) or 48 hr (IF). C) PCA plot for RNA-seq samples with mESC (blue) and NE (pink) treated with 24 hours iASCL1 (darker) or not (lighter). Simultaneously, samples were either treated with DMSO (circle) or 10 nM TSA (triangle). D) Examples of previously identified NE-specific ASCL1-respsonsive genes that become ASCL1-responsive in mESC with ASCL1 + TSA treatment. D) *Nkx6-1*; D’) *Disp2*; D’’) *Plxna2*. Adjusted p-values from DESeq2 differential gene expression analysis. ns p > 0.05, ** p < 0.01, *** p < 0.001 E) Heatmap of z-scored RNA-seq counts for genes that were identified as NE-specific ASCL1 targets in Fig. 1H–I and have associated ASCL1 binding site (n = 208). Expression data from mESC: control, iASCL1 only, TSA only, iASCL1+TSA. E’) genes of the 208 in the heatmap that are significantly upregulated in the comparisons listed: iASCL1+TSA versus control (n = 116); TSA only versus control (n = 102); iASCL1+TSA versus TSA only (n = 13). F) Heatmap of z-scored RNA-seq counts for genes that are significantly upregulated in TSA+iASCL1 versus TSA only (n = 458). F’) Gene ontology analysis for all genes in heatmap (F). Top 6 GO terms are listed. “Regulation of neurogenesis” highlighted with black arrow. G) IF images of H3K27ac (magenta) and TUBB3-positive neurites (green) after 48 hours ±ASCL1 and ±TSA in mESC or NE. DAPI counterstain (blue). Scale bar = 100 µm.

TSA treatment altered the transcriptome of both mESCs and NE cells with and without iASCL1 (Fig. 5C and S5B–F) (Table S5). To assess whether TSA could alter the responsiveness of NE-specific ASCL1 target genes in mESCs, we focused on this previously defined set of genes (Fig. 2D). Several of these genes have established roles in neurogenesis, including *Nkx6-1*, *Disp2*, and *Plxna2* (Fig. 5D–D’’), and all three of these were now upregulated by iASCL1 in mESCs when co-treated with TSA (Fig. 5D–D’’). More broadly, combined iASCL1+TSA treatment drove the highest expression of NE-specific ASCL1 target genes in mESCs compared to iASCL1 or TSA alone, with 116 genes significantly upregulated relative to the control in this condition (Fig. 5E–E’).

However, TSA alone had a substantial independent effect on the mESC transcriptome compared to untreated cells, upregulating 4,924 genes including 102 of the previously identified NE-specific iASCL1 targets (Fig. S5D and Fig. 5E’), suggesting that many of these changes reflect opportunistic activation by other factors in response to broadly increased acetylation rather than directed ASCL1 activity.

Beyond analysis of the previously identified NE-specific targets described above, broader transcriptional analysis of differentially expressed genes in mESCs revealed many additional genes that responded to combined iASCL1+TSA treatment relative to TSA alone (n = 458) (Fig. 5F and S5F). GO analysis of these genes returned terms including “placenta development” and “mesenchyme development” as the top hits, alongside “regulation of neurogenesis” (Fig. 5F’), indicating that co-treatment activates a largely incoherent and generally non-neuronal transcriptional programme. Consistent with this, co-treatment with iASCL1 and TSA failed to produce TUBB3-positive neurons within 48 hours (Fig. 5G). This is likely because global changes to the acetylation landscape caused by TSA treatment expose additional sites primed for ASCL1 binding at non-neuronal targets, generating conflicting alternative transcriptional programmes. Taken together, these results show that globally increasing histone acetylation can sensitise individual NE-specific target genes to ASCL1 in mESCs but is insufficient to redirect ASCL1 activity towards a productive neuronal differentiation programme and away from divergent pathways.

### Homeodomain motifs are enriched at ASCL1 binding sites

We next asked whether we could use more targeted approaches to engage a neuronal ASCL1 programme in mESCs, which do not have an epigenetic landscape that favours directed neuronal differentiation. We reasoned that, despite the availability of open neuronal gene-associated ASCL1 binding sites, ASCL1 activation of neuronal target genes may be limited in mESCs because essential co-binding TFs that normally guide ASCL1 activity to these sites in an NE context are absent from mESCs. To identify cofactors capable of enhancing ASCL1 activity at genomic sites associated with neuronal genes, we examined potential cofactor motif enrichment in NE-specific ASCL1 binding sites compared against the set of universally bound sites (Fig. 6A), prioritising motifs associated with transcription factors found at higher levels in NE relative to mESCs before ASCL1 activation (Fig. 6A). We confirmed that these motifs were also enriched in the 653 sites associated with NE-specific iASCL1 target genes (Fig. S6B). This analysis identified Sox (HMG) and Homeodomain-containing TFs as the dominant motif families at NE-specific ASCL1 binding sites. In contrast, mESC-specific ASCL1 binding sites were enriched for motifs associated with nuclear receptor, zinc finger and AP2 TFs (Fig. S6A), indicating distinct cofactor environments in mESCs and NE.

**Figure 6.**
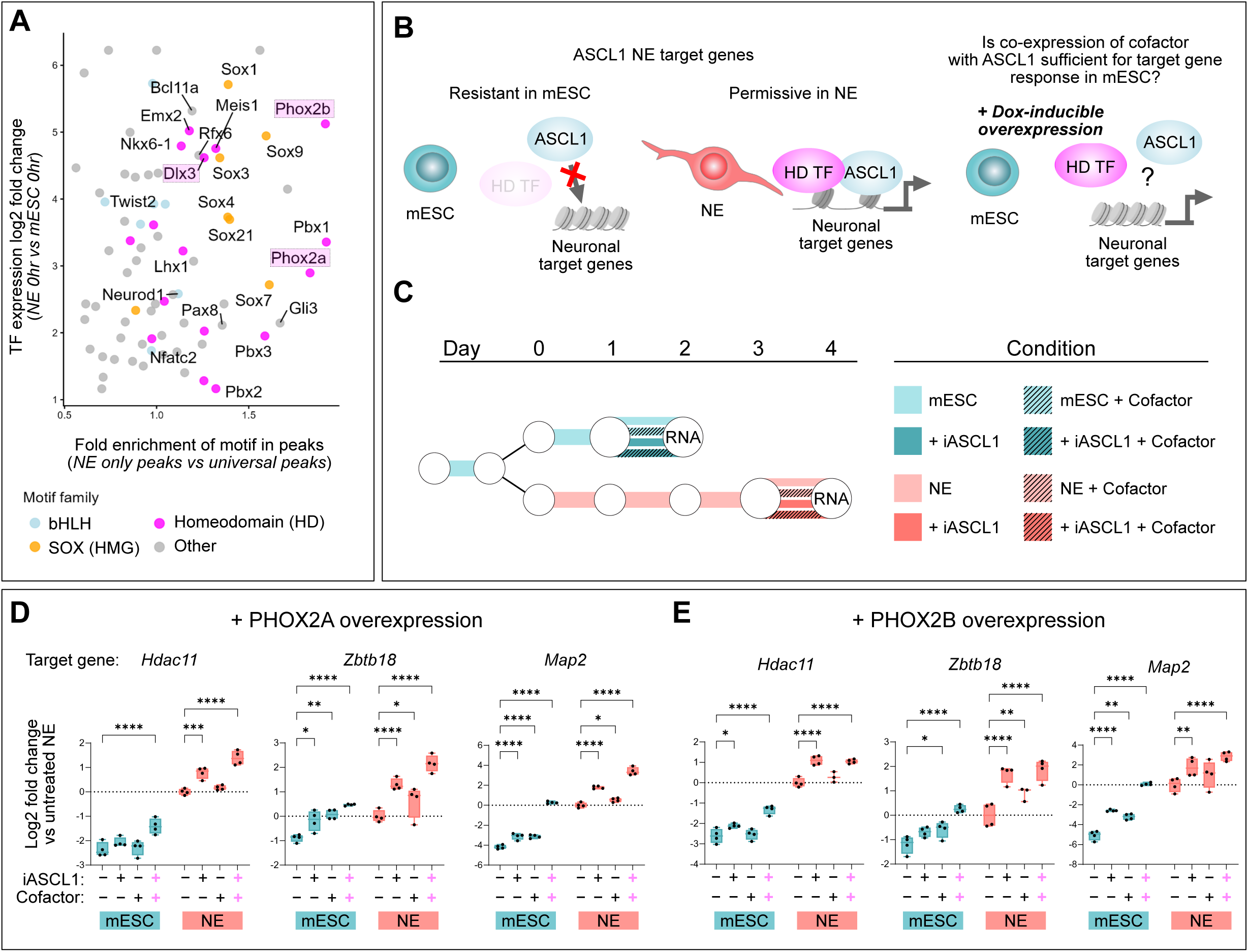
Homeodomain transcription factor motifs are enriched in NE-specific ASCL1 binding sites and co-expression with ASCL1 enables activation of previously NE-specific target genes. A) Plot of TFs scored by fold enrichment of motif in NE-specific peaks (x) against log fold change in expression comparing NE versus mESC (No iASCL1) from RNA-seq (Fig. 1). Coloured by TF family. Phox2b, Phox2a, and Dlx3 are highlighted as shortlisted candidates. B) Schematic for putative model of cooperativity between ASCL1 and Homeodomain (HD) TFs. C) Schematic for culture and treatment of mESC (blue) and NE (pink) for 24 hrs: control (light), +iASCL1 only (dark), +cofactor (Dox induction) only, +iASCL1+cofactor D–E) qPCR of NE-specific ASCL1 DEGs associated with ASCL1 binding sites that contain HD motif (*Hdac11* and *Zbtb18*) and NE-specific marker of neurogenesis (*Map2*). mESC (blue) and NE (pink). Dox used to induce overexpression of cofactors, either PHOX2A (D) or PHOX2B (E). Control, iASCL1 alone, Dox alone, or iASCL1 and Dox. N = 4 experiments, each performed in technical duplicate. Two-way ANOVA with Dunnett’s correction for multiple comparisons to the control for each cell type.

SOX4 and SOX11 are known to act downstream of bHLH proteins in neuronal development,^77,78^ while SOX1 is expressed in neuroectoderm. SOX2 is expressed in both pluripotency and neural stem cells, so is unlikely to be the limiting factor for ASCL1-mediated neuronal differentiation in mESCs. Top candidate homeodomain cofactors identified in our analysis included PHOX2B, PHOX2A, and DLX3 (Fig. 6A and Fig. S6B). DLX3 has been reported to potentiate ASCL1 reprogramming of fibroblasts.^41^ We therefore decided to focus on relatively unexplored homeodomain-containing transcription factors as potential ASCL1 cofactors in potentiating context-dependent coherent neuronal programming. We also included LMX1A as a representative homeodomain (HD) TF with a known role in neural development^79^ that was not enriched in the motif analysis to test whether any functional effects we saw were more generalisable to other homeodomain-containing TFs.

### Homeodomain TFs potentiate ASCL1-driven neuronal differentiation in mESCs

To investigate whether ASCL1 and HD TFs act cooperatively to drive neurogenesis, we used a doxycycline-inducible piggyBac expression system^80^ to conditionally overexpress four candidate cofactors alone or together with iASCL1 in mESCs and NE cells (Fig. S6C–E). We first asked whether co-expression could activate previously unresponsive ASCL1 neuronal target genes in mESCs after 24 hours, focusing on *Zbtb18* and *Hdac11*, which are both associated with a proximal ASCL1 binding site containing a homeodomain TF motif. Co-expression of PHOX2A, PHOX2B, or DLX3 with ASCL1 increased expression of both genes in NE cells and further potentiated the effect of ASCL1 alone (Fig. 6D–E and Fig. S6F), while LMX1A showed less consistent effects across targets (Fig. S6G). Critically, although neither *Zbtb18* nor *Hdac11* responds to iASCL1 alone in mESCs, co-expression with PHOX2A, PHOX2B, or DLX3 was sufficient to upregulate both genes. Co-activation also increased expression of *Map2*, a downstream neuronal differentiation marker and NE-specific iASCL1 target gene, in mESCs (Fig. 6D–E and Fig. S6F–G). These results demonstrate that selected HD TFs can enhance ASCL1 responsiveness at individual neuronal target genes in otherwise resistant mESCs.

To investigate how co-expression of these cofactors with ASCL1 affects transcriptional targets genome-wide in mESCs, we performed bulk RNA-seq on mESCs overexpressing PHOX2B with or without iASCL1 after 24 hours (Fig. 7A–B and Fig. S7A) (Table S6). Firstly, we saw that the number of upregulated genes in iASCL1+PHOX2B mESCs (n = 757) was greater than that observed in iASCL1 alone (n = 480) or PHOX2B alone (n = 240), indicating a compound effect of both factors together (Fig. S7A). We then looked at how the iASCL1+PHOX2B upregulated genes behaved across conditions (Fig. 7A). We saw that these genes fit broadly into two clusters: those that have highest expression in the iASCL1+PHOX2B-treated mESCs (Fig. 7A, Cluster 1) and those that have comparable expression in mESCs treated with either iASCL1 alone or iASCL1+PHOX2B (Fig. 7A, Cluster 2). This indicates that the combined activity of these two transcription factors creates a synergistic new set of target genes that are otherwise silent. GO analysis on both Cluster 1 (Fig. 7A’) and Cluster 2 (Fig. 7A’’) identified “regulation of neurogenesis”, but “axon guidance” was only enriched in Cluster 1. This could indicate that the genes upregulated by combined activity of iASCL1 and PHOX2B are associated with a more mature or coherent neuronal transcriptional programme.

**Figure 7.**
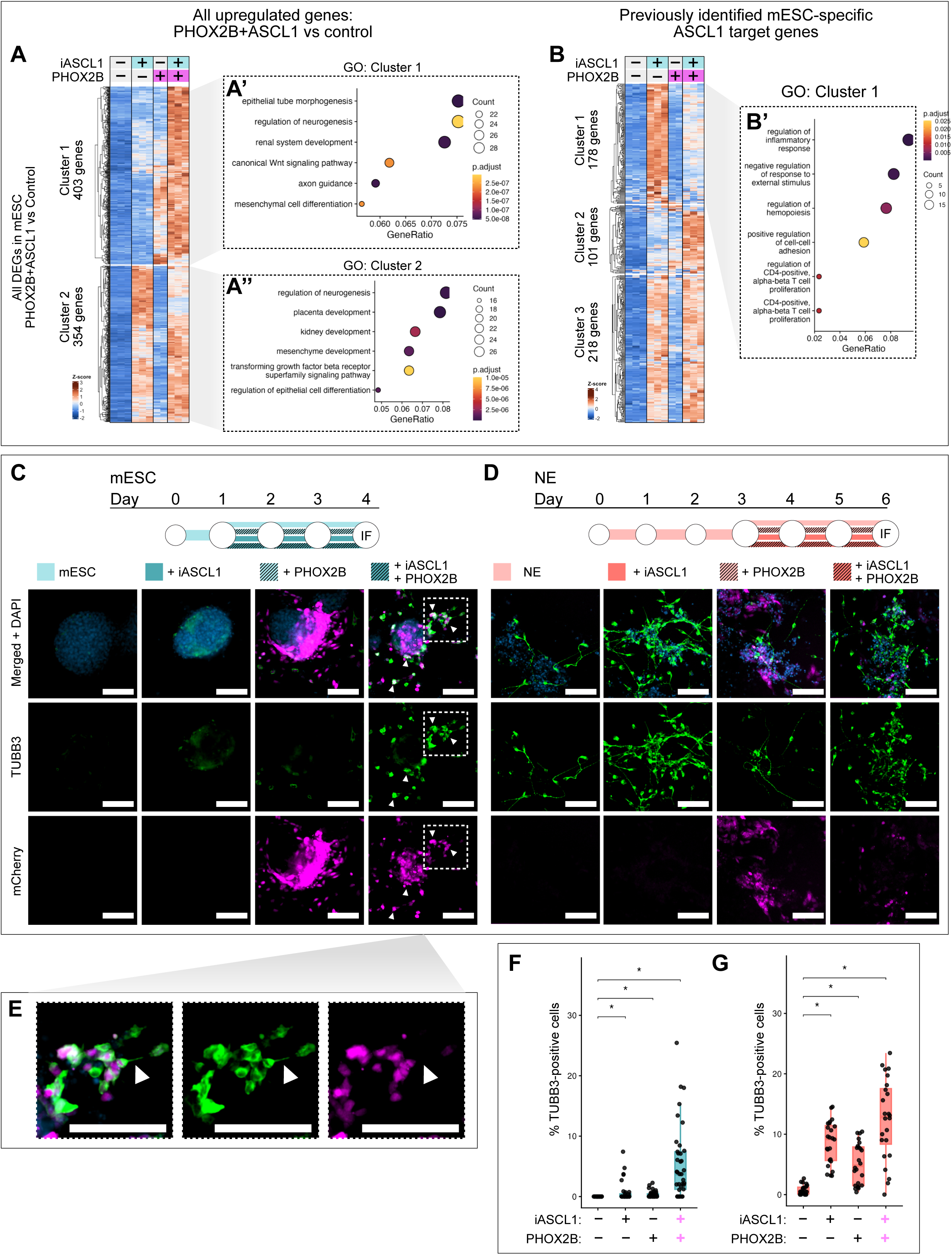
Genome-wide redirection of ASCL1 transcriptional output with PHOX2B co-expression. A) Heatmap of z-scored RNA-seq counts for genes that are significantly upregulated by iASCL1+PHOX2B relative to untreated control mESCs (n = 757). Genes are clustered into 2 groups: Cluster 1 shows highest enrichment in iASCL1+PHOX2B cells (n = 403); Cluster 2 shows comparable expression in iASCL1 alone and iASCL1+PHOX2B. A’) GO analysis for genes in Cluster 1. A’’) GO analysis for genes in Cluster 2. Top 6 terms are shown. B) Heatmap of z-scored RNA-seq counts for genes that were previously identified as mESC-specific iASCL1 targets (Fig. 1H) (n = 497). Genes are clustered using k means = 3. B’) GO analysis of genes in Cluster 1. Top 6 terms are shown. C–D) Schematic for culture and treatment of mESC (C, blue) and NE (D, pink) before collection for immunofluorescence (IF) imaging. Cells were either untreated, iASCL1 only, PHOX2B only, or iASCL1+PHOX2B. TUBB3 (green), mCherry (magenta), DAPI (blue). mCherry and DAPI LUTs are adjusted separately for mESC and NE, but TUBB3 is consistent across conditions. Scale bar = 100 µm. Representative images from 2 experiments each done in duplicate. E) Magnified inset from Fig. 7C. White arrow indicates neurite-like outgrowth. Scale bar = 100 µm. F–G) Quantification of TUBB3-positive cells in cofactor differentiation assay in mESC and NE untreated, with ASCL1 alone, Dox alone, or ASCL1 and Dox. Each dot represents an image taken from a randomised position in the well. Statistics are performed using N = 4 for 4 wells, from 2 independent passages. Wilcoxon pairwise test with Benjamini–Hochberg post-hoc correction. * p < 0.05

ASCL1 activation in ESCs results in upregulation of divergent gene programmes that may actively interfere with coherent neuronal programming. We therefore asked whether previously identified ASCL1 target genes upregulated specifically in mESCs (Fig. 1H–I) displayed an altered response to ASCL1 when it was co-expressed with PHOX2B. (Only 497 of the previously defined 601 mESC-specific iASCL1 targets were detected in this RNA-seq experiment due to differences in library preparation and sequencing depth.) A subset of these genes (178 out of 497) showed less activation in mESCs when iASCL1 activation was combined with PHOX2B activation (Fig. 7B) than with iASCL1 alone, and these genes were associated with terms such as “regulation of hemopoiesis” and “T cell proliferation” (Fig. 7B’). Therefore, PHOX2B may work with ASCL1 both to facilitate the activation of neuronal target genes and also to suppress the activation of non-neuronal targets that interferes with successful differentiation in mESCs.

Finally, we assessed whether these transcriptional changes we observed in mESCs in response to iASCL1 and HD TFs translated into neuronal differentiation, assayed by TUBB3 expression and cell morphology after a longer 72-hour period compared to earlier analysis of more immediate iASCL1 effects (Fig. 1–5). In NE, which already express some endogenous PHOX2B and PHOX2A (Fig. 6A), additional ectopic PHOX2B and PHOX2A both induced a modest increase in neurogenesis compared to control. However, neither PHOX2B nor PHOX2A co-expression with iASCL1 significantly increased the proportion of TUBB3-positive cells relative to iASCL1 alone (PHOX2B: 3.0% versus 7.7%, p > 0.05; PHOX2A: 9.2% versus 12.9%, p > 0.05) (Fig. 7D and 7G and Fig. S7C–D). DLX3 and LMX1A did not increase TUBB3-positive cell numbers in NE even when expressed alone or with iASCL1 (Fig. S7E–F), highlighting differences in the intrinsic neurogenic capacity of individual HD TFs.

In mESCs, iASCL1 activation alone produced only sparse, diffuse TUBB3 expression after 3 days (Fig. 7C). PHOX2B and PHOX2A expression did not result in TUBB3-positive cells in the absence of iASCL1 (Fig. 7C and 7F; Fig. S7B and S7D). However, strikingly, co-expression of PHOX2B or PHOX2A with iASCL1 produced TUBB3-positive cells with immature neuronal morphology (Fig. 7C and 7E–F; S7B and S7D). DLX3 and LMX1A failed to generate TUBB3-positive cells in mESCs with or without iASCL1 (Fig. S7E–F), indicating that the ability to confer neuronal competence on ASCL1 in mESCs is specific to PHOX2A and PHOX2B and not a general feature of homeodomain-containing TFs.

## DISCUSSION

Together, these data support a model in which ASCL1 is not an intrinsically neuron-specific pioneer factor, but a highly context-dependent transcription factor whose activity is constrained by both chromatin state and cofactor availability. Cell-intrinsic changes during the exit from pluripotency are required for competence for ASCL1-mediated neuronal differentiation and to channel ASCL1 activity away from activating divergent programmes. However, this neuronal competence can be increased by altering the global chromatin landscape or by providing additional cofactor TFs that enhance neuronal target gene activation.

Many studies have used mESCs (and human iPSCs) for forward programming and TF-mediated neuronal differentiation. However, it is important to note that many of them incorporate embryoid body formation, or some other priming step.^7,9^ Vainorius et al. use ASCL1 to drive neuronal differentiation from mESCs, but with two important differences: firstly, they cultured their mESCs in serum-LIF, in which ESCs are heterogeneous and display epigenetic and transcriptional priming of many lineage commitment genes^57,81^; secondly, cells were plated on Matrigel and differentiated in N2B27 after doxycycline induction of ASCL1 expression. Thus, these cells have likely already transitioned through formative pluripotency and are competent for ASCL1-mediated differentiation. Casey et al. profiled genome-wide binding of ASCL1 in mESCs grown on feeder cells, and observed a similar binding profile of ASCL1 to our mESC data, but did not investigate competence for neuronal differentiation or the expression of non-neuronal genes.^46^ We find that while failing to upregulate a coherent neuronal programme, pluripotent mESCs are nevertheless fully competent to respond transcriptionally to ectopic ASCL1, but in this context non-neuronal heterologous gene programmes dominate; exit from pluripotency is required to allow an appropriate ASCL1-specific developmental programme to be engaged. This underscores the importance of chromatin remodelling to restrict the cell identity during the early stages of embryonic development.^76,82,83^

Despite the fact that pluripotent cells have the ability to differentiate into any embryonic cell type and are characterised by having globally low levels of repressive heterochromatin,^12,73^ it is perhaps surprising to find that chromatin accessibility played a major role in determining ASCL1 binding in ESCs. Rather than a model with widespread chromatin accessibility in pluripotent cells offering a “blank canvas” for highly permissive binding of transcription factors, with an epigenetic environment progressively refined during development, our data point to complex but distinct landscapes. Unlike accessible sites bound by ASCL1 in NEs, sites accessible to ASCL1 binding in ESCs are generally not associated with neuronal genes. Moreover, in the context of gene activation in ESCs, EpiLCs and NEs, we see a limited ability of ASCL1 to act as a pioneer opening chromatin. Recent data in human iPSC differentiation to neurons, has shown ASCL1 displays different modes of activity at different sites across the genome, sometimes acting alone and sometimes recruiting SWI/SNF chromatin remodelling complexes to act as a non-classical pioneer factor.^35^ Our study adds to this model, showing that ASCL1 acts as both a pioneer and a non-pioneer TF to varying degrees across the genome and between cell types. Moreover, there must be additional layers of regulation that determine whether a given binding site is accessible to ASCL1-mediated pioneering. Differential availability of chromatin remodelling complexes across the developmental time course may determine whether a given site can be pioneered by ASCL1, as has been described for SWI/SNF subunit switching in neuronal differentiation.^35,84^

Accessibility, though clearly important, is not the only factor that determines context-dependent ASCL1 binding and gene activation. Many ASCL1 binding sites near neuroectoderm-specific target genes were marked by H3K27ac in neural progenitors but not in mESCs or EpiLCs, even when accessibility is comparable at these sites across cell types. This is especially interesting in light of the prevailing view of mESCs being broadly unrestricted in terms of their epigenetic landscape and developmental potential. Instead, ASCL1 binding sites that are activated in NE are maintained in inactive chromatin in mESC and EpiLC and are not poised or primed. This likely reflects the fact that they are not the first regulatory elements recruited *in vivo* in the exit from pluripotency, as ASCL1 itself is expressed at later stages of embryo development.^85,86^ Consistent with this, global elevation of histone acetylation using an HDAC inhibitor renders many previously unresponsive neuronal genes inducible by ASCL1 in mESCs, highlighting the importance of chromatin priming in establishing developmental competence. Even transient treatment with HDAC inhibition can cause significant changes in genome architecture in mESCs,^87^ and our transcriptomic data indicates that TSA treatment creates many additional non-neuronal ASCL1 target genes. HDAC inhibitors have been used to potentiate ASCL1-mediated reprogramming in the adult retina,^88^ but it is possible that additional epigenetic constraints in these tissues limit the ectopic effects of induced ASCL1 activity.

To identify more targeted approaches to modulating ASCL1 activity in mESCs, we identified candidate co-factors by identifying specific motifs for neuroepithelial transcription factors found near to ASCL1 binding sites in NE target genes. We found that co-expression of ASCL1 with candidate homeobox TFs in mESCs allows activation of many previously NE-specific neuronal genes while simultaneously inactivating genes associated with divergent pathways. Interaction between HD TFs and bHLH factors, including ASCL1, has been previously shown in the diversification of cell types in the developing retina^89^ cranial neural crest cells.^90^ PHOX2A and PHOX2B have been implicated in gene regulatory networks with ASCL1 in the development of the central nervous system,^91–93^ sympathetic neurons,^94,95^ and in neuroblastoma.^96,97^ ASCL1 expression typically precedes and is required for PHOX2 factor expression in these cases, so it is notable that we show simultaneous expression of both factors potentiated the direct programming capacity of ASCL1 in mESCs, where PHOX2 factors cannot be upregulated by ASCL1 activation directly. This suggests that the sequential logic of endogenous development can be bypassed by simultaneous cofactor delivery.

It is also possible that cell type-specific activity of ASCL1 is the result of differential post-translational modification. Phosphorylation, for example, has been shown to modulate the activity of various bHLH TFs in development and reprogramming.^98–102^ However, our recent work showed that a phosphorylation-resistant ASCL1 mutant was not sufficient to drive neuronal differentiation in mESCs,^10^ so this is unlikely to explain why only NE cells undergo ASCL1-mediated neuronal differentiation in these experiments.

In summary, we demonstrate that despite their ability to differentiate into any embryonic cell type, the competence of ES cells to respond to the lineage-determining transcription factor ASCL1 is limited by a chromatin landscape in which non-neuronal enhancers are favourable for ASCL1 binding, which instead directs ASCL1 activity to divergent genes and pathways. Our findings illustrate the multifaceted controls that limit reprogramming in different cellular contexts and provide a framework for improving directed differentiation and reprogramming approaches by targeting competence rather than transcription factor expression alone.

## SUPPLEMENTARY FIGURE LEGENDS

**Figure S1 – Related to Figure 1.**
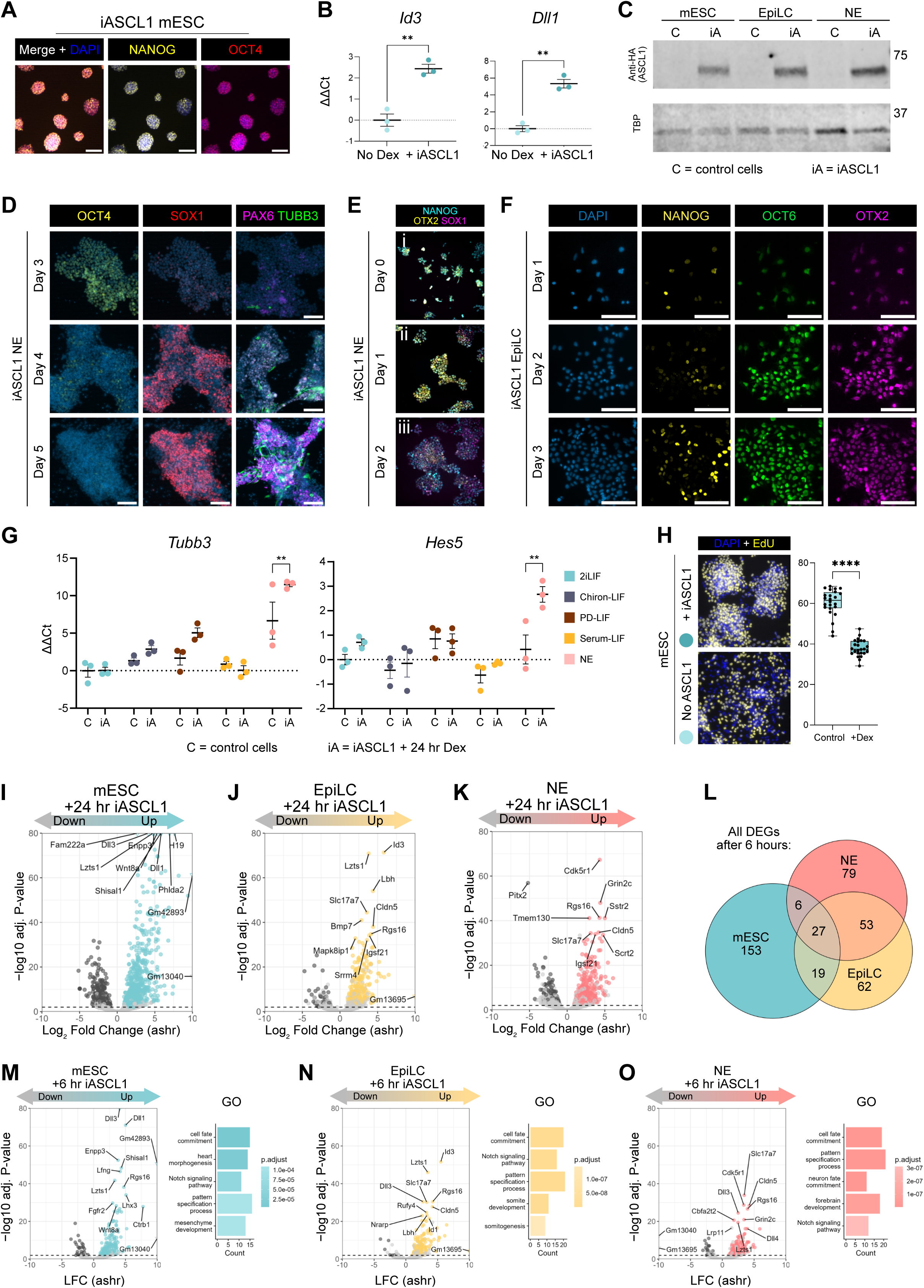
A) iASCL1 mESC express OCT4 (green) and NANOG (red) in 2iLIF. Scale bar = 50 µm. B) qPCR of ASCL1 target genes ± 24h Dex in iASCL1 mESC in 2iLIF. N = 3 experiments. ** p < 0.01, t-test. C) Western blot to show HA-ASCL1-GR is specific to iASCL1 and levels are consistent across differentiation conditions (mESC, EpiLC, and NE). Representative of 2 western blots. Molecular weight markers are indicated (right). D) IF of OCT4 (yellow), SOX1 (red) and PAX6 (magenta) in iASCL1 differentiated into NE conditions on day 3-5. Spontaneous neuronal differentiation from day 4 onwards indicated by TUBB3 (green). Scale bar = 100 µm. Representative of 2 experiments. E) IF of NANOG (blue), OTX2 (yellow) and SOX1 (magenta), day 0-2 of NE differentiation. F) IF for naïve pluripotency marker, NANOG (yellow), and formative markers, OCT6 (green) and OTX2 (magenta) in iASCL1 cells on days 1-3 of EpiLC differentiation. Scale bar = 100 µm. G) qPCR of ASCL1 target genes in 2iLIF variant media. N = 3 independent experiments. Two-way ANOVA with Dunnett’s post-hoc pairwise comparisons comparing control versus +iASCL1 for 24 hours. H) EdU-positive mESCs (yellow) with or without 24 hours of iASCL1 activity. Quantification of positive cells from 8 images from 3 independent experiments. **** p < 0.0001, t-test. I–K) Volcano plots to show upregulated genes in mESC (I), EpiLC (J) and NE (K) after 24 hours of iASCL1 relative to untreated cells in each of: mESC (blue), EpiLC (yellow), NE (pink). Note: some genes appear above threshold for significance but are coloured grey and excluded from analysis based on comparison between untreated cells and Dex-treated control cells with no iASCL1-GR transgene to control for Dex-specific effects. L) Venn diagram to show overlap in DEGs after 6 hours. M–O) Volcano plots and GO analysis on upregulated genes in mESC (M), EpiLC (N) and NE (O) after 24 hours of iASCL1 relative to untreated cells in each of: mESC (blue), EpiLC (yellow), NE (pink).

**Figure S2 – Related to Figure 2.**
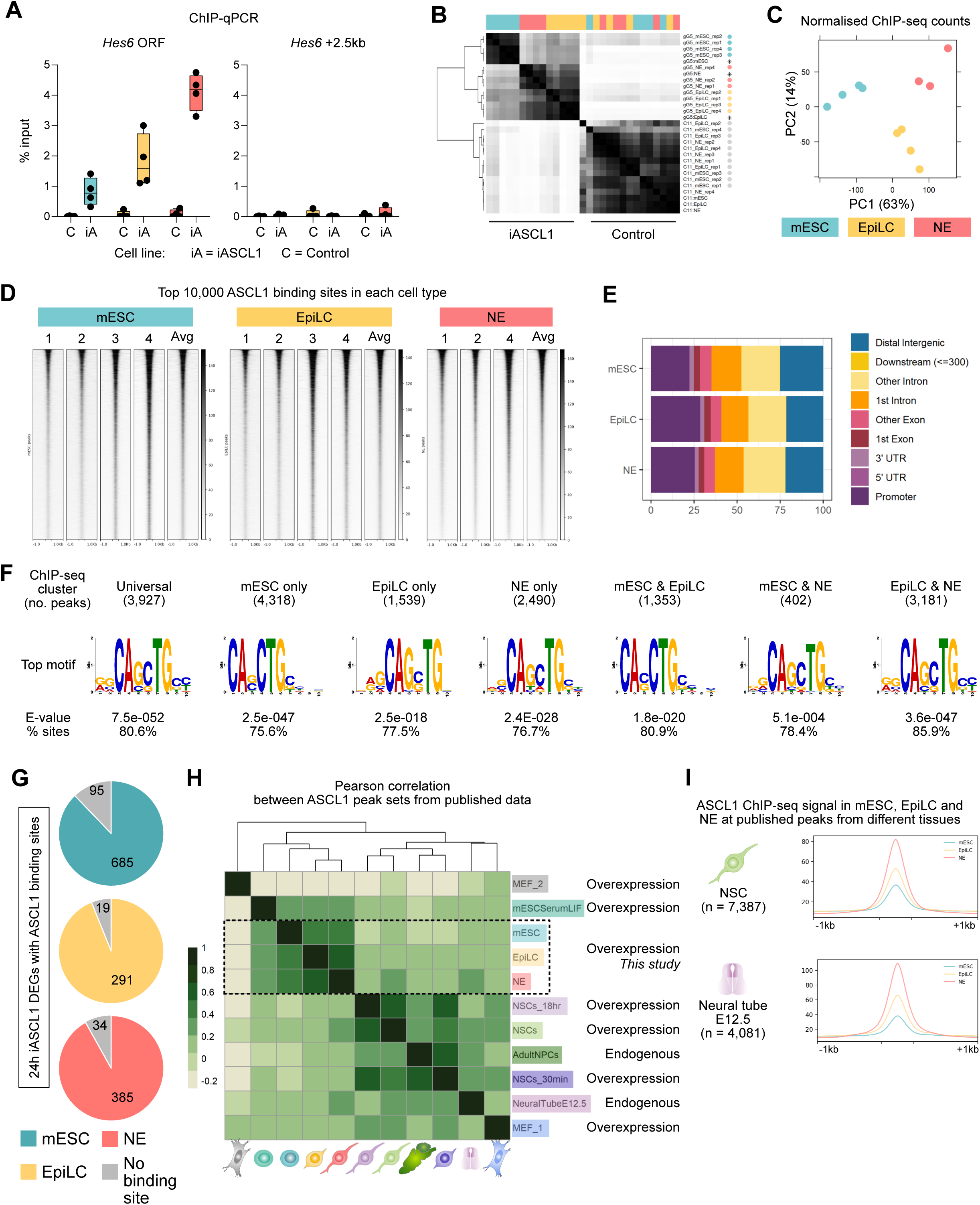
A) ChIP-qPCR to validate the anti-HA ASCL1 ChIP-seq. N = 4 independent experiments. B) Correlation matrix of ChIP-seq peak sets, including all iASCL1 samples and control cells that have no iASCL1-GR transgene. C) PCA plot to show sample distances after normalisation (using RLE method and reads in peaks as library size). D) Heatmap across replicates after normalisation at the top 10,000 peaks in the consensus set. E) Annotation of top 10,000 ASCL1 binding sites in each cell type. Promoter regions are defined as ±1kb from TSS. F) Top motif discovered by STREME in each of the clusters. E-value and percentage of target sites that contain the motif are reported next to each motif. G) Number of differentially expressed genes (DEGs) in mESC (blue), EpiLC (yellow) and NE (pink) that have a binding site within ±100 kb. DEGs with no proximal ASCL1 binding site are grey. H) Correlation matrix for publicly available ASCL1 ChIP-seq datasets, including mESC, EpiLC and NE from this study. Full GEO accession numbers are available in the Methods section. I) Profiles of ChIP-seq signal in mESC, EpiLC, and NE from this study at peaks from neural stem cells (NSC),^43^ and embryonic neural tube.^72^

**Figure S3 – Related to Fig.3.**
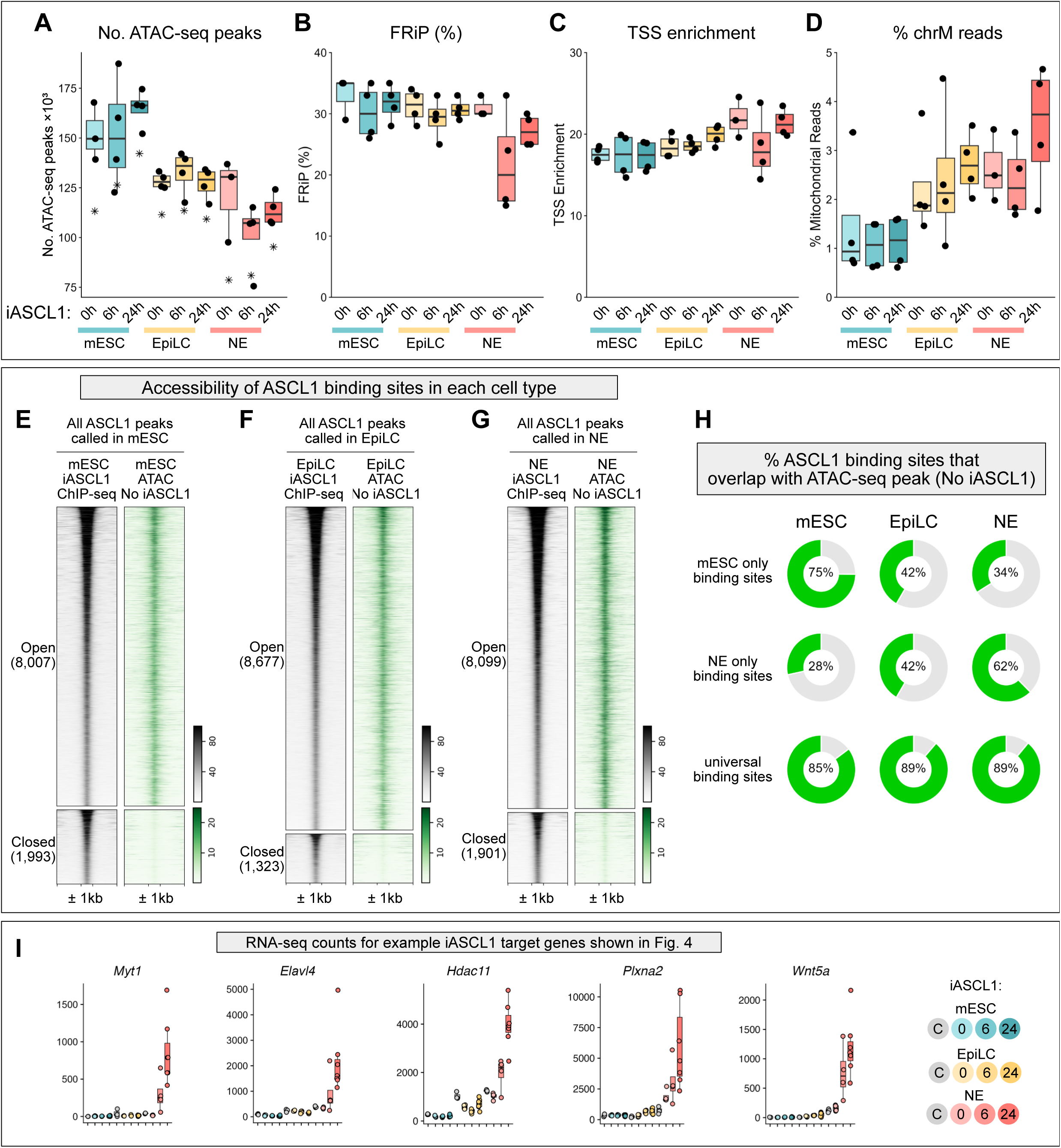
A) Number of peaks called in each sample. Asterisk corresponds to the per-sample consensus peak set, comprising peaks called in at least 3 of the 4 replicates. B) Fraction of reads in peaks (FRiP) for all samples. C) TSS enrichment is the ratio of reads the align to TSS of known genes over the reads in 1000 bp regions flanking the TSS. D) Percentage of reads that align to mitochondrial DNA per sample. E) iASCL1 ChIP-seq (black) and No iASCL1 ATAC-seq (green) data in mESC at all 10,000 ASCL1 binding sites detected in mESC, split by whether they are open (top) or closed (bottom) in the No iASCL1 ATAC-seq data. F) iASCL1 ChIP-seq (black) and No iASCL1 ATAC-seq (green) data in EpiLC at all 10,000 ASCL1 binding sites detected in EpiLC, split by whether they are open (top) or closed (bottom) in the No iASCL1 ATAC-seq data. G) iASCL1 ChIP-seq (black) and No iASCL1 ATAC-seq (green) data in NE at all 10,000 iASCL1 binding sites detected in NE, split by whether they are open (top) or closed (bottom) in the No iASCL1 ATAC-seq data. H) Donut plots for percentage of iASCL1 ChIP-seq peaks in cell type-specific clusters (Fig. 2A) that overlap with an ATAC-seq peak (green) or not (grey) in each cell type. I) RNA-seq counts for NE-specific ASCL1-responsive DEGs highlight in Fig.3. As per RNA-seq experiment, samples were either treated with 0, 6, or 24 hours of iASCL1 and compared to a control line with no iASCL1 transgene for each cell type.

**Figure S4 – Related to Fig.4.**
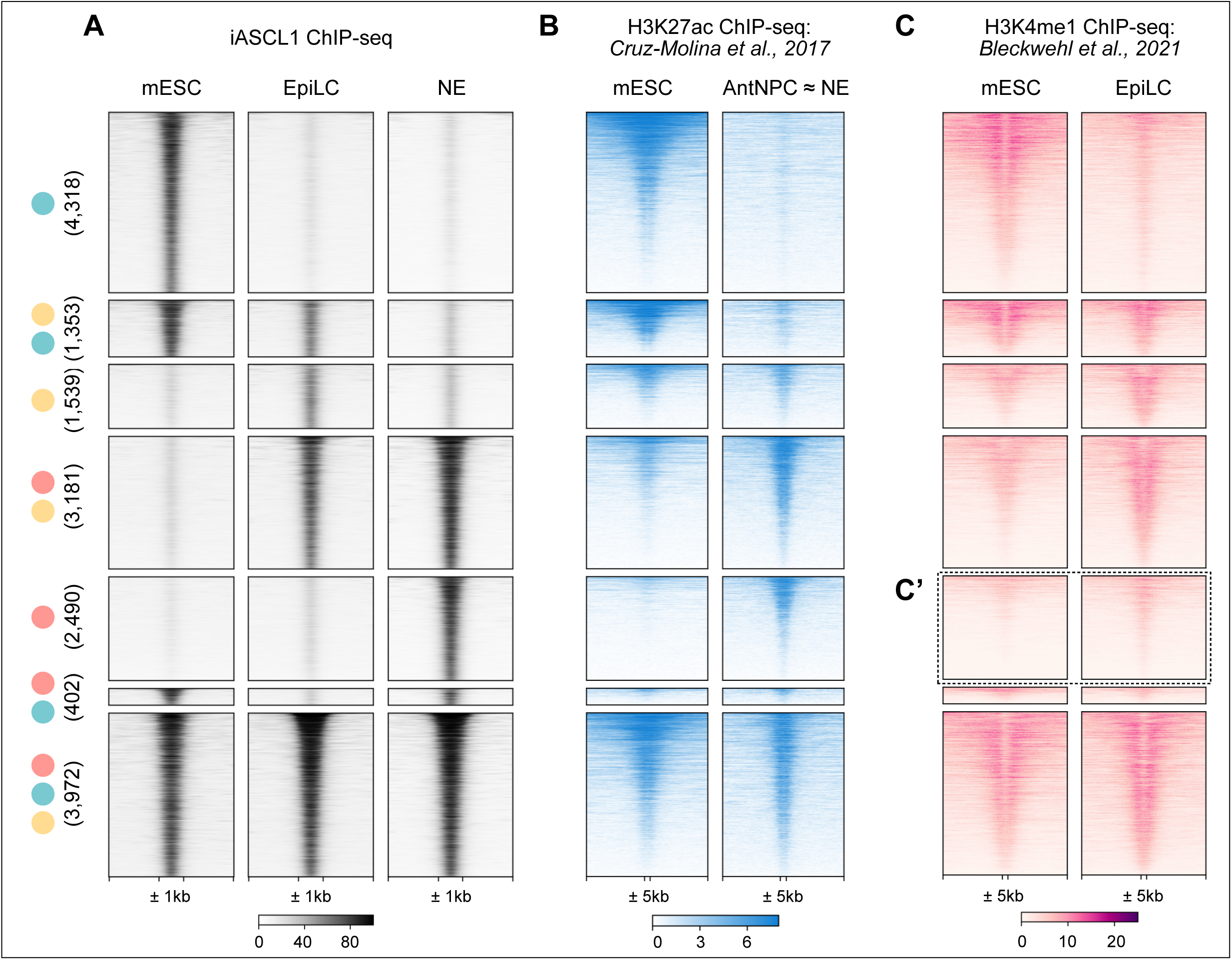
A) ASCL1 ChIP-seq as in Fig. 2 and 3. B) H3K27ac signal in mESC and AntNPC from Cruz-Molina et al., 2017^20^ plotted across ASCL1 binding sites (this study) split by cell type-specific clusters (Fig. 2A). Centred around ASCL1 binding site (500 bp) ± 5 kb. C) H3K4me1 signal in mESC and EpiLC from Bleckwehl et al., 2021^76^ plotted across ASCL1 binding sites (this study) split by cell type-specific clusters (Fig. 2A). Centred around ASCL1 binding site (500 bp) ± 5 kb. C’) highlights the NE-specific iASCL1 binding sites.

**Figure S5 – Related to Figure 5.**
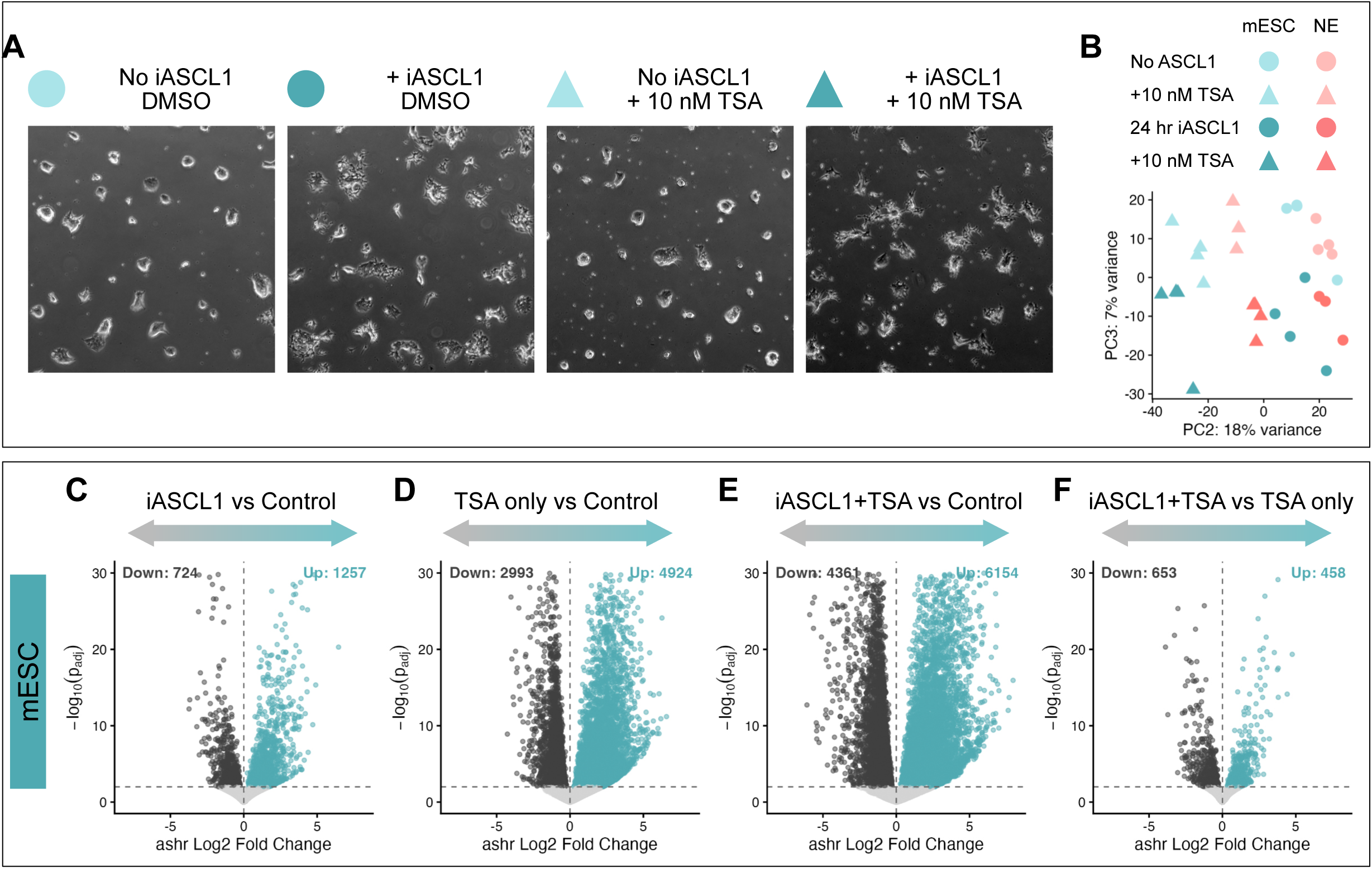
A) Phase contrast to show morphology and density of mESC treated with nothing, +iASCL1 only, 10 nM TSA only, or +iASCL1 and +TSA. B) PCA plot of PC2 and PC3 from RNA-seq. C–F) Volcano plot to show differentially expressed genes in mESC in specific comparisons: C) iASCL1 versus control; D) TSA only versus control; E) iASCL1+TSA versus control; F) iASCL1+TSA versus TSA only. Significance thresholds: absolute log fold change > 0 and adjusted p-value < 0.01. ashr shrunken log fold changes are used for plotting.^103^

**Figure S6 – Related to Figure 6.**
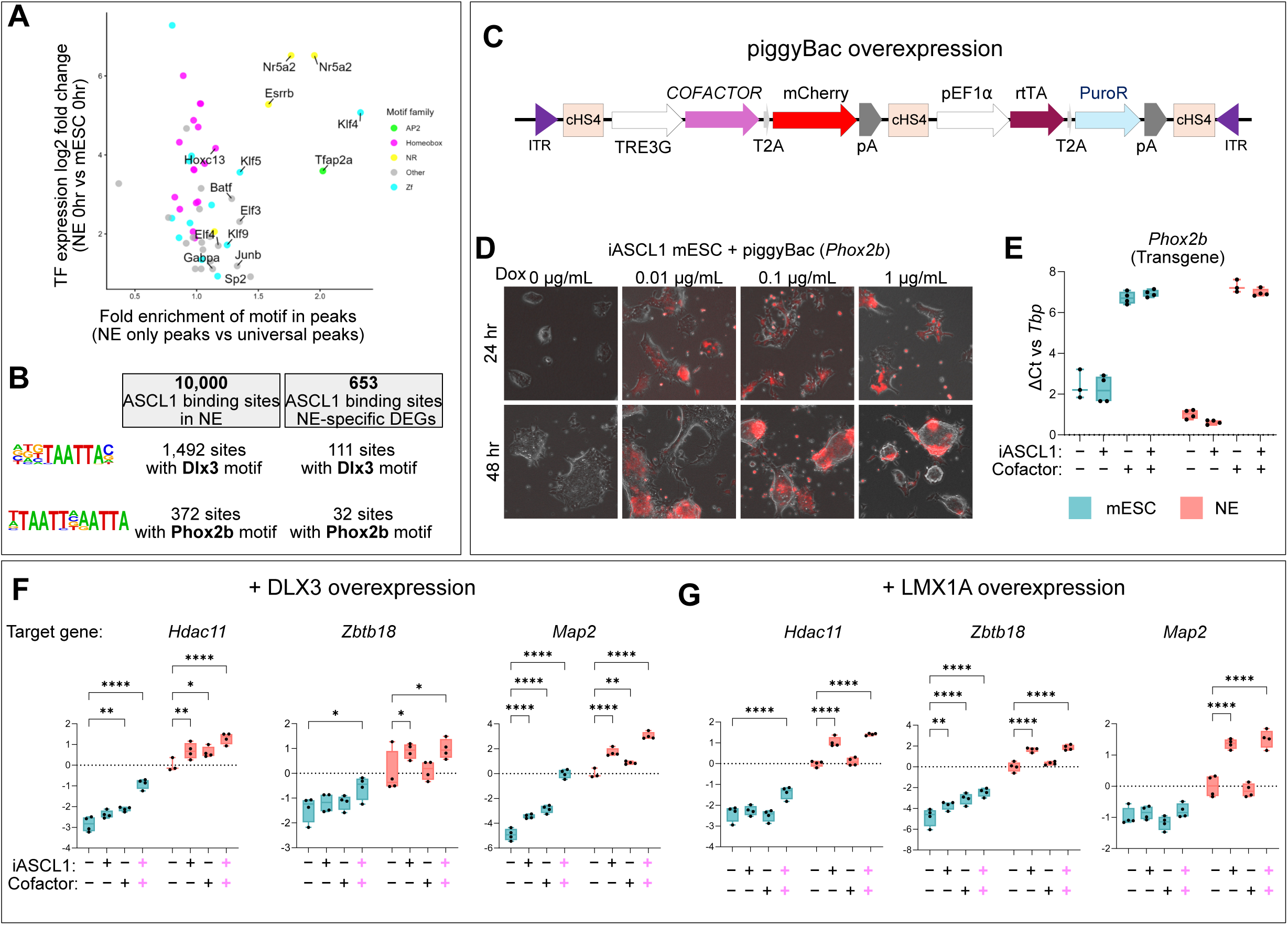
A) Plot of TFs scored by fold enrichment of motif in mESC-specific peaks (x) against log fold change in expression in mESC versus NE at t = 0. Coloured by TF family. B) Number of ASCL1 binding sites that contain a DLX3 or PHOX2B motif. C) piggyBac construct used for Dox-inducible overexpression. ITR, inverted terminal repeat; cHS4, chicken hypersensitive site 4 insulator; TRE3G, third-generation tetracycline-responsive promoter; COFACTOR is where candidate cofactors were cloned in; T2A, self-cleaving peptide; mCherry; pA, polyadenylation signal; pEF1⍺ promoter; rtTA Tet-On activator; PuroR, puromycin resistance gene. D) Titration of Dox treatment on pool of antibiotic-resistant cells. mCherry (red) indicates induction of the transgene. E) qPCR of *Phox2b* transgene induction relative to *Tbp* housekeeping. N = 4 experiments F–G) qPCR of NE-specific ASCL1 DEGs associated with ASCL1 binding sites that contain HD motif (*Hdac11* and *Zbtb18*) or *Map2*. Dox used to induce overexpression of either DLX3 (F) or LMX1A (G). Cells were untreated or treated with either iASCL1 alone, Dox alone, or iASCL1 and Dox. N = 4 experiments, each performed in technical duplicate. Two-way ANOVA with Dunnett’s correction for multiple comparisons to each cell type’s respective control.

**Figure S7 – Related to Figure 7.**
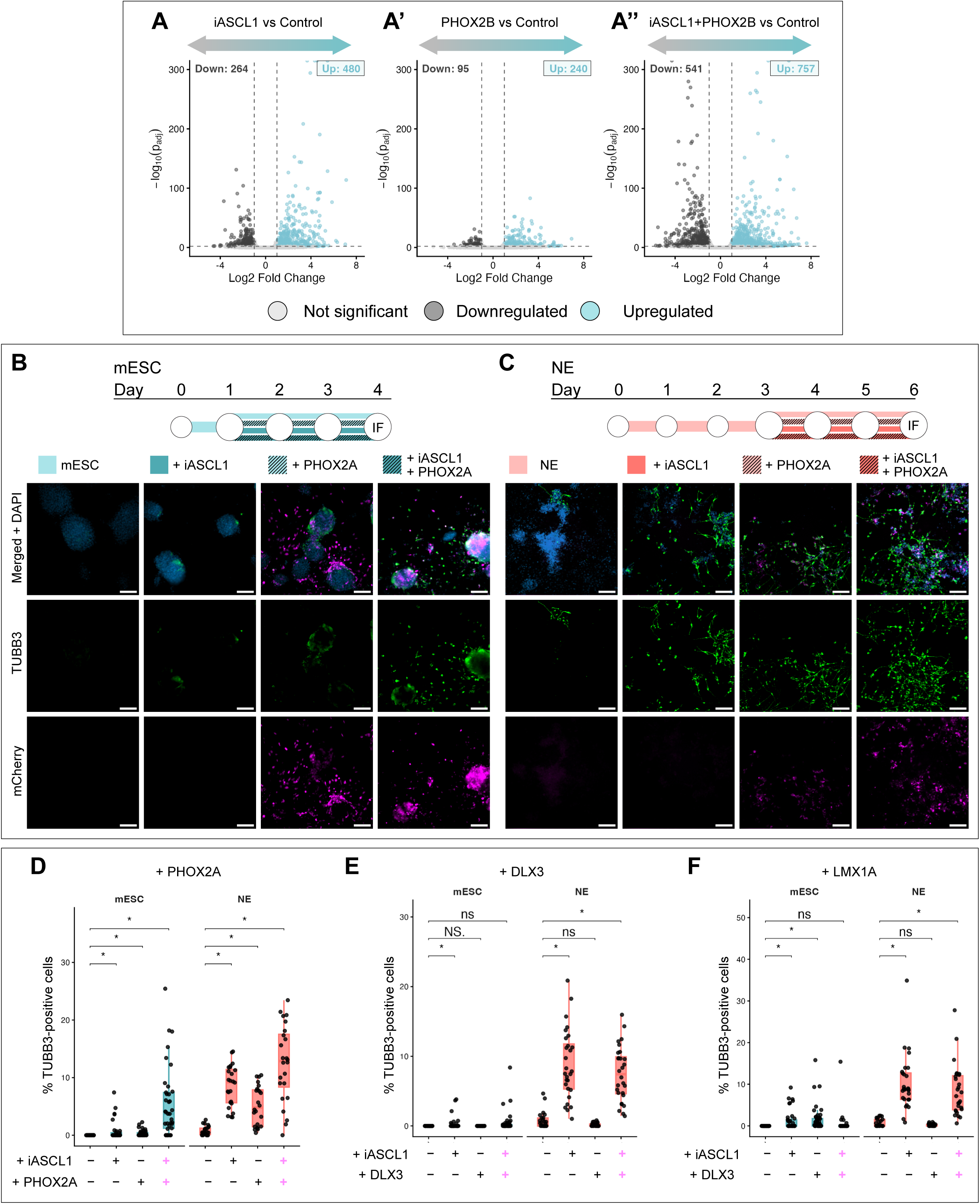
A) Volcano plots for specific comparisons in mESC from RNA-seq (Fig. 7A–B): iASCL1 versus control; PHOX2B only versus control; iASCL1+PHOX2B versus control. Significance thresholds: absolute log fold change > 1 and adjusted p-value < 0.01. ashr shrunken log fold changes are used for plotting.^103^ B–C) Schematic for culture and treatment of mESC (B, blue) and NE (C, pink) before collection for immunofluorescence (IF) imaging. Cells were either untreated, iASCL1 only, PHOX2A only, or iASCL1+PHOX2A. TUBB3 (green), mCherry (magenta), DAPI (blue). mCherry and DAPI LUTs are adjusted separately for mESC and NE, but TUBB3 is consistent across conditions. Scale bar = 100 µm. Representative images from 2 experiments each done in duplicate. D–F) Quantification of TUBB3-positive cells in cofactor differentiation assay in mESC and NE. Cofactors are PHOX2A (D), DLX3 (E) and LMX1A (F). Each dot represents an image taken from a randomised position in the well. Statistics are performed using N = 4 for 4 wells, from 2 independent passages. Wilcoxon pairwise test with Benjamini–Hochberg post-hoc correction. * p < 0.05

## METHODS

### Cell culture

Mouse embryonic stem cells (mESCs; ES-E14TG2a, ATCC #CRL-1821) were maintained as described previously.^104^ Ascl1-GR-HA mESCs (iASCL1; R. Azzarelli, unpublished) were cultured in 2i/LIF medium: N2B27 basal medium (1:1 Neurobasal [Gibco, 21103-049] / DMEM-F12 [Gibco, D6421], N2 [in-house, Cambridge Stem Cell Institute], B-27 [Gibco, 17504044], 2 mM L-glutamine, 50 µM β-mercaptoethanol) supplemented with 100 U/mL leukaemia inhibitory factor (LIF; in-house), 3 µM CHIR-99021 (ABCR, AB253776) and 1 µM PD0325901 (ABCR, AB253775).

Cells were passaged every 2–3 days using Accutase (300 × g, 3 min) and seeded at 1.5 × 10⁴ cells/cm² on 0.1% gelatin-coated 6-well plates. Stocks were cryopreserved at 1 × 10⁶ cells/mL in 50% 2i/LIF / 40% KSR / 10% DMSO. Mycoplasma testing was performed every 3–6 months.

For lineage transitions, mESCs were plated on fibronectin (16.7 µg/mL) to generate epiblast-like cells (EpiLC) in N2B27 + FGF2 (12 ng/mL) + Activin A (20 ng/mL),^3,5^ or on laminin (10 µg/mL) for primitive neuroectoderm (NE) differentiation in N2B27 + 0.1% sodium bicarbonate + 0.11% BSA + 20 µg/mL insulin.^104^

A gene fragment encoding N-terminal HA-tagged full-length ASCL1 fused to the glucocorticoid receptor ligand-binding domain (GR-LBD), followed by an IRES-GFP cassette, was synthesised with flanking attB1/attB2 sites (Twist Bioscience). Gateway LR Clonase II (Thermo Fisher) was used to insert the construct into a Rosa26-targeting destination vector (SP202) carrying appropriate homology arms.^105^ E14 mESCs were co-transfected with the targeting vector, Rosa26-directed sgRNAs, and a Cas9 expression vector using the 4D Nucleofector (Lonza). Correctly targeted clones were isolated by FACS for constitutive GFP expression, expanded, and verified by PCR across both homology arms for monoallelic (heterozygous) insertion. All experiments used a single validated clonal line.

In iASCL1 mESCs, ASCL1 activity is post-translationally controlled: the HA-ASCL1-GR fusion protein is constitutively expressed but held inactive by HSP90 chaperone binding until displacement by 5 µM dexamethasone (Dex).^106^

### Cofactor overexpression mESCs

PiggyBac-based, doxycycline (Dox)-inducible overexpression vectors were assembled from two published plasmids: the TRE3G-driven expression cassette was PCR-amplified (CloneAmp, Takara, 639298) from pPB-Ins-TRE3Gp-KRAB-dCas9-ecDHFR-IRES-GFP-EF1Ap-Puro (Addgene #183410), and the rtTA-Neo cassette was amplified from pPB-Ins-U6p-sgRNAentry-EF1Ap-TetOn3G-IRES-Neo (Addgene #183411). Both were cloned into a custom piggyBac-based vector (VB240515-1632gka, VectorBuilder) via EcoRV/PacI digestion and In-Fusion Snap Assembly (Takara, 638948). The dCas9-KRAB-U6-sgRNA module was excised with NotI/AsiSI and replaced with synthesised transgene sequences (PHOX2A, PHOX2B, LMX1A, and DLX3: Twist Bioscience; mCherry: VectorBuilder). Gel purification used the Monarch kit (NEB, T1120); plasmids were propagated in Stellar competent cells (Takara, 636763), purified with QIAprep Miniprep or Midiprep kits (Qiagen), and sequence-verified by Nanopore sequencing (Plasmidsaurus).

For PHOX2A, PHOX2B, LMX1A, and DLX3 overexpression lines, mESCs were co-transfected with PB-Dox-Cofactor plasmid and Super PiggyBac Transposase vector (Stratech, PB210PA-1-SBI; 500 ng) to a total of 2 µg DNA, complexed with 3.5 µL Lipofectamine 2000 (Invitrogen, 11668027) in 400 µL Opti-MEM™ and incubated on cells for 4 h at 37°C. Medium was replaced with 2i/LIF, then selection in 0.5 µg/mL puromycin (2i/LIF) initiated 48–72 h post-transfection and maintained until colonies were pooled. Overexpression efficiency was confirmed by RT-qPCR after 1 µM Dox induction.

### Immunofluorescence microscopy

Cells were fixed in 4% PFA (ThermoFisher, J19943-K2) for 20 min at room temperature, permeabilised in 0.2% Triton-X100 (Sigma, X100)/PBS for 15 min, and blocked in 5% serum / 0.01% Triton-X100 / PBS for 1 h. Primary antibodies (PBS + 2% serum + 0.01% Triton-X100) were applied overnight at 4°C; secondary antibodies for 1 h at room temperature. Washes used PBS-T (PBS + 0.01% Triton-X100); DAPI (1 µg/mL, Abcam, ab228549) was added during the third wash (20 min). Images were acquired on a Leica DMI6000 at 10× or 20×.

S-phase fraction was used as a proliferation proxy. Cells were treated with 5 µM Dex for 24 h; then 10 µM EdU (final; Click-iT EdU Kit, Alexa Fluor 488, Thermo Fisher, C10637) was added for 1 h before fixation. EdU was detected per manufacturer’s instructions. Note: iASCL1 mESC constitutive GFP was negligible relative to AlexaFluor-488-labelled targets, so the 488 channel was used without interference.

### Image analysis

Nuclear intensities were quantified in CellProfiler v4.0.^107^ Nuclei were segmented from DAPI using the IdentifyPrimaryObjects module (Otsu two-class adaptive thresholding; smoothing factors determined empirically) and filtered by size. Channel intensities within each nuclear object were measured; binary positivity thresholds were set empirically. Percentages of positive/negative nuclei were calculated from ≥10 images across 2–3 biological replicates.

For TUBB3 quantification (Fig. 6), nuclear ROIs were expanded by 5 pixels to capture peri-nuclear signal, which was used as a proxy for total TUBB3 (individual cell-level quantification was not feasible in these morphologically complex monolayers). Intensities were normalised to the control sample mean; statistical comparisons used per-replicate means (N = 4, two separate passages, each performed in duplicate), not individual nucleus counts.

### RT-qPCR

RNA was extracted with the RNeasy Mini Kit (Qiagen, 74106) and stored at −80°C. cDNA was synthesised with the QuantiTect Reverse Transcription Kit (Qiagen, 205314), including DNase treatment; cDNA was diluted 1:5–1:10 prior to qPCR. Reactions (10 µL, 384-well) were run on a QuantStudio 7 Flex or ViiA 7 Real-Time PCR System. Primer efficiency was validated by serial dilution of a pooled mESC/EpiLC/NE cDNA reference; primers with an efficiency outside 85–115% were excluded. Quantification used the modified ΔΔCt method incorporating primer efficiency.^108^ TBP and GAPDH served as housekeeping genes. Expression ratios were log-transformed prior to statistical testing to satisfy normality.

### Bulk RNA-seq

RNA was extracted as above and treated with the DNA-free DNA Removal Kit (Invitrogen, AM1906). Poly(A) RNA was enriched using the NEBNext Poly(A) mRNA Magnetic Isolation Module (E7490); strand-specific libraries were prepared with the NEBNext Ultra II Directional RNA Library Prep Kit (E7760) and NEBNext Multiplex Oligos (E6440). Sequencing was performed as 50 bp paired-end reads on NovaSeq 6000 or NovaSeq X (Illumina).

For cofactor overexpression experiments, RNA-seq was performed by Plasmidsaurus (Illumina) using their in-house analysis pipeline; raw counts matrices were used for downstream analysis. Cells (∼10⁵ per condition) were collected in 50 µL DNA/RNA Shield (Zymo, R1100-50) after Accutase dissociation (300 × g, 3 min).

### Computational analysis of RNA-seq

Raw BCL files were converted to FASTQ with bcl2fastq (Illumina). Adapter trimming and quality filtering (Phred ≥ 33) used Trim Galore; trimmed reads were pseudo-aligned to the mouse genome (mm39) with Kallisto.^109^

Kallisto output was imported into R with tximport (DESeq2 package).^110^ Genes with fewer than 10 counts in fewer than 3 samples were removed. Differential expression analysis used DESeq2; DEGs (Fig. 1) were defined as |log₂FC| > 0.585 (≥1.5-fold) and adjusted p < 0.01. Fold change thresholds were > 0 for HDACi RNA-seq (Fig. 5) and > 1 for cofactor experiment (Fig. 7) to account for different variance within these datasets. Variance-stabilised transformation was used for PCA, distance matrix, and heatmap visualisation.^110^ ashr was used for shrinkage estimation in volcano plots.^103^ Gene Ontology enrichment used enrichGO (clusterProfiler).^111^

### ChIP-seq

ChIP-seq was performed as described^112^ with modifications. Cells were crosslinked in 1% formaldehyde (10 min, room temperature), quenched with 125 mM glycine, washed twice in ice-cold PBS, and pelleted (2,000 × g, 4 min). Pellets were flash-frozen on dry ice and stored at −80°C. Sequential lysis used: LB1 (50 mM HEPES-KOH pH 7.5, 140 mM NaCl, 1 mM EDTA, 10% glycerol, 0.5% NP-40, 0.25% Triton-X100; 10 min, ice, 2,000 × g), LB2 (10 mM Tris-HCl pH 8, 200 mM NaCl, 1 mM EDTA, 0.5 mM EGTA; 10 min, ice, 2,000 × g), and LB3 (10 mM Tris-HCl pH 8, 100 mM NaCl, 1 mM EDTA, 0.5 mM EGTA, 0.1% Na-deoxycholate, 0.5% N-lauroylsarcosine). All buffers contained cOmplete protease inhibitor (Sigma, 4693159001).

Chromatin was sheared using a Bioruptor Plus (Diagenode, 4°C; 30 s on/off, 16 cycles in 2 rounds of 8) to yield 100–600 bp fragments (verified by agarose gel after heat reversal). Chromatin concentration was determined by A280 on a NanoDrop (2 µL in 98 µL 0.1 M NaOH). Per IP, 100 µg chromatin was diluted to 300 µL in LB3 + 1% Triton-X100 and clarified (16,000 × g, 10 min); 10% was retained as input. IP was performed overnight with anti-HA antibody (ab9110) pre-loaded on BSA-blocked Protein G Dynabeads (Invitrogen). Beads were washed 10× in RIPA buffer (50 mM HEPES-KOH, 500 mM LiCl, 1 mM EDTA, 1% Igepal CA-630, 0.7% Na-deoxycholate) and once in TBS (20 mM Tris-HCl pH 7.6, 150 mM NaCl). DNA was eluted overnight at 65°C in 200 µL elution buffer (50 mM Tris-HCl pH 7.6, 10 mM EDTA, 1% SDS), which simultaneously reversed crosslinks.

Eluted DNA was diluted 1:1 in TE, then purified by sequential RNase A digestion (40 µg/mL, 37°C, 30 min), proteinase K digestion (400 µg/mL, 55°C, 2 h), phenol-chloroform-isoamyl alcohol phase separation in MaXtract HD tubes (Qiagen), and ethanol precipitation (200 mM NaCl, −20°C) with 1 µL GlycoBlue (20 µg/µL, Thermo Fisher, AM9515) as co-precipitant. Purified DNA was resuspended in 20 µL 10 mM Tris-HCl pH 8. Enrichment was validated by qPCR at known ASCL1 target loci (percentage input from 10% fraction). Libraries were prepared using ThruPLEX DNA-Seq (Takara, R400675) at the JCBC NGS Facility and sequenced on NovaSeq 6000 at 15 nM equimolar pooling.

### Computational analysis of ChIP-seq data

Reads were trimmed with Trim Galore (Phred ≥ 33) and aligned to a composite mm39/dm6 index with Bowtie2. *Drosophila* reads (spike-in) were discarded as normalisation by spike-in was not applied; spike-ins were included as a precautionary measure, but RLE normalisation was sufficient for comparison between conditions. mm39-aligned reads were used for all downstream analysis. Peaks were called with MACS3^113^ using matched input as background control. Specificity was confirmed by the near-absence of peaks in control (no-transgene) samples; any such peaks were excluded from the consensus set. The consensus peak set comprised peaks present in ≥2 of 4 replicates per condition. Reads-in-peaks normalisation used RLE (DESeq2). Peaks were annotated to transcription start sites within 100 kb using AnnotationHub^114^ and Ensembl *Mus musculus* v108.

Published ENCODE mouse ASCL1 ChIP-seq datasets were downloaded via accession numbers (Materials Table) and converted from mm10 to mm39 using UCSC liftOver. ^115^ Inter-dataset Pearson correlation and Jaccard similarity were assessed from a binary peak-presence matrix in DiffBind.

### Motif analysis

De novo motif discovery was performed using MEME-ChIP (MEME Suite, web server),^116^ which applies STREME for improved sensitivity at centrally enriched motifs,^117^ and independently with HOMER^118^ (-size 150, -mask) for both de novo and known-motif enrichment.

### ATAC-seq

ATAC-seq was performed using the Omni-ATAC protocol.^119,120^ To synchronise multi-timepoint collection, cells were seeded on sequential days. Cells were dissociated with Accutase, counted with Trypan Blue, and 5 × 10⁴ cells resuspended in ice-cold PBS were pelleted and lysed in 50 µL RSB++ (10 mM Tris-HCl pH 7.4, 10 mM NaCl, 3 mM MgCl₂, 0.1% NP-40, 0.1% Tween-20, 0.01% digitonin) for 3 min on ice. Nuclei were washed with 1 mL RSB+ (RSB++ without NP-40/digitonin), pelleted (500 × g, 10 min, 4°C), and resuspended in 50 µL Tn5 transposition mix. Tagmentation was performed at 37°C for 30 min with 1,000 rpm mixing, then cleaned with Zymo DNA Clean & Concentrator (D4014) and stored at −20°C.

Tagmented DNA was PCR-amplified with indexed i5/i7 primers, purified with the Qiagen MinElute kit (eluted in 20 µL EB), and size-selected with a dual AMPure XP bead (#A63880) strategy: (i) 0.55× beads added to remove large fragments (supernatant retained), then (ii) 1.5× beads to capture the library; beads washed 2× in 80% ethanol and eluted in 20 µL EB. Fragment size distribution was confirmed by Bioanalyzer HS D1000 (Agilent, #5067-5584). Libraries were pooled equimolarly and sequenced on NovaSeq X 25B (PE100), targeting ∼9 × 10⁷ reads/sample.

### Computational analysis of ATAC-seq

FASTQ files were generated with bcl2fastq2. A custom bash pipeline based on nf-core ATAC-seq^121,122^ and published best practices^123^ was applied. Quality control used FastQC followed by Trim Galore (default paired-end parameters). Reads were aligned to mm39 with Bowtie2^124^ (--very-sensitive-local, --no-mixed, --no-discordant; maximum insert size 2,000 bp). SAMtools^125^ was used to index, sort, and assess alignment statistics; mitochondrial reads were removed. Duplicates were marked with Picard and removed; insert size metrics were collected with Picard CollectInsertSizeMetrics. Peaks were called with MACS3^113^ in BAMPE mode (effective genome size 2,468,088,461).

Downstream analysis used R v4.5.0 in RStudio. Peaks overlapping ENCODE blacklisted regions^126^ or unplaced/random contigs were removed. DiffBind generated a consensus peak set (peaks present in ≥3 of 4 replicates per condition); reads-in-peaks were TMM-normalised with DiffBind. Bigwig files were produced with bamCoverage (deepTools) ^127^ using TMM scale factors; per-condition averages used bamCompare. Visualisation used IGV^128^ and deepTools heatmaps.

## ACKNOWLEDGEMENTS

We would like to thank the Cambridge Stem Cell Institute Genomics Facility at the Jeffrey Cheah Biomedical Centre for all of their help with library preparation and sequencing; the Cambridge Stem Cell Institute Imaging Core Facility; Prof. Brian Hendrich and members of his lab group for their assistance with ChIP-seq and ATAC-seq; William Beckman, Lidiya Mykhaylechko, Sarah Gillen, Frances Connor, and all members of the Philpott lab for helpful discussions throughout; and Dr Mekayla Storer, Prof. Jason Carroll and Dr Srinjan Basu for their useful guidance.

This work was supported by a Wellcome Trust Investigator award (212253/Z/18/Z) to (A.P., J.L.-B., R.D., J.-P.N.-B.); J.L.-B. is funded by a Wellcome Trust studentship (218481/Z/19/Z); R.A. is supported by core funding from UCL School of Pharmacy and by the Royal Society grant (RGS\R2\242202).

## AUTHOR CONTRIBUTIONS

J.L.-B. performed most of the experimental work and bioinformatic analyses. R.A. generated the iASCL1 mESC line with contribution from R.D. J.L.-B., R.D. and J.-P.N.-B generated the piggyBac overexpression system for cofactor experiments. R.A. and J.L.-B. designed the experiments. J.L.-B. prepared the figures with input from R.A and A.P. R.A. and A.P. supervised the project and obtained funding. J.L.-B., R.A. and A.P wrote the manuscript with input from R.D. and J.-P.N.-B.

## SUPPLEMENTAL MATERIAL

Table S1 (differentially expressed genes per condition, relating to Fig. 1 and S1), Table S2 (GO analysis, relating to Fig. 1 and S1), Table S3 (ASCL1 peaks linked to proximal genes, relating to Fig. 2), Table S4 (653 NE-specific ASCL1 binding sites, relating to Fig. 3 and Fig. 4), Table S5 (differentially expressed genes with HDACi treatment, relating to Fig. 5), Table S6 (differentially expressed genes with PHOX2B and ASCL1, relating to Fig. 7)

## RESOURCE AVAILABILITY

### Lead contact

Further information and requests for resources and reagents should be directed to and will be fulfilled by the lead contacts, Dr Roberta Azzarelli or Prof. Anna Philpott.

### Materials availability

All unique/stable reagents and cell lines generated in this study are available from the lead contact with a completed materials transfer agreement.

### Data and code availability

Bulk RNA-seq, ChIP-seq and ATAC-seq data have been deposited at GEO (GSE327328) and are publicly available as of the date of publication. Published ASCL1 ChIP-seq data were accessed from: GSM2576033, GSM1175113, E-MTAB-2384, E-MTAB-2384, GSE43916, GSE43916, GSM1347006, GSE89209. Microscopy data reported in this paper will be shared by the lead contact upon request. All original code is available via GitHub (github.com/Philpott-lab/ascl1_competence). Any additional information required to reanalyse the data reported in this work is available from the lead contact upon request.

## DECLARATION OF GENERATIVE AI AND AI-ASSISTED TECHNOLOGIES IN THE WRITING PROCESS

During the preparation of this work the author(s) used Claude v1.2773.0 for proofreading of the manuscript and to assess R and bash code reproducibility. After using this tool/service, the author(s) reviewed and edited the content as needed and take(s) full responsibility for the content of the published article.

